# Inhibition of ITK differentiates GVT and GVHD in allo-HSCT

**DOI:** 10.1101/2020.07.15.204693

**Authors:** Mahinbanu Mammadli, Weishan Huang, Rebecca Harris, Aisha Sultana, Ying Cheng, Wei Tong, Jeffery Pu, Teresa Gentile, Jessica Henty-Ridilla, Shanti Dsouza, Qi Yang, Avery August, Alaji Bah, Mobin Karimi

## Abstract

Allogeneic hematopoietic stem cell transplantation is a life-saving treatment for many malignant and nonmalignant diseases. Donor T cells contained within the graft prevent tumor recurrence *via* graft-versus-tumor (GVT) effects, however, also cause graft-versus-host disease (GVHD). Novel treatment strategies are therefore needed to allow maintenance of GVT while suppressing GVHD. Here we show using murine models, that targeting IL-2-inducible T cell kinase (ITK) in donor T cells reduces GVHD while preserving the beneficial GVT effects. Donor T cells from *Itk_-/-_* mice exhibit significantly reduced production of inflammatory cytokines and migration to GVHD target organs such as liver and small intestine, while maintaining GVT efficacy against primary B-ALL tumors. *Itk_-/-_* T cells exhibited reduced expression of IRF4 and decreased JAK/STAT signaling activity, but preserved cytotoxicity, which was accompanied by upregulation of Eomesodermin (Eomes), which was necessary for GVT function. A novel peptide inhibitor ITK signaling is also able to prevent GVHD. This novel peptide inhibitor also reduced cytokine production in mice and human T cells. Altogether, our data suggest that inhibiting ITK could be a therapeutic strategy to reduce GVHD while preserving the beneficial GVT effects following allo-HSCT treatment.

**Key Points:**

- Inhibiting ITK by a novel peptide significantly reduces GVHD but retains GVT.
- ITK deficient donor T cells exhibit minimal GVHD, but maintain GVT activity.
- ITK deficient donor T cells exhibit significantly reduced production of inflammatory cytokines and migration to GVHD target organs.
- Eomes is required for GVT effect.

## Introduction

Allo-reactive donor T cells present in the graft are required for donor stem cell engraftment and are essential for anti-tumor activity (graft-versus-tumor: GVT)^1, 2^, but the same donor T cells also cause significant tissue damage to the host known as graft-versus-host disease (GVHD)^3^. This remains the most significant obstacle to the broader application of allogeneic hematopoietic stem cell transplantation (allo-HSCT), a life-saving treatment for many malignant and nonmalignant diseases. Standard immunosuppressive therapy for GVHD is often therapeutically sub-optimal and predisposes patients to opportunistic infections such as Cytomegalovirus (CMV) and relapse of the underlying malignancy^4, 5^. Thus, specific signaling pathways that can be targeted to allow the effects of GVT to occur while inhibiting GVHD need to be identified.

The Tec family nonreceptor tyrosine kinase, Interleukin-2-inducible T cell kinase (ITK), regulates activation of T cells downstream of T cell receptor (TCR) and has been shown to be involved in the activation of intracellular calcium signaling and MAPK pathways, and polarization of actin cytoskeleton, supporting an integral role for ITK in T cell activation and function^6, 7^. ITK regulates T cell signaling by participating in a multimolecular proximal signaling complex, which includes the adaptor SH2 domain-containing leukocyte protein of 76 kDa (SLP76)^8, 9^. In particular, ITK interacts with tyrosine residue 145 (Y145) of SLP76 when phosphorylated, and a Y145F mutation of SLP76 leads to defective TCR-mediated ITK activation^10^. Interaction between SLP76 and ITK not only influences signals leading to cytokine-production by T cell populations, but also regulates the development of distinct innate type cytokine-producing T cell populations in the thymus^11, 12, 13^, referred to as innate memory phenotype (IMP) T cells.

Since activation, expansion, cytokine production, and migration of alloreactive donor T cells to target organs are hallmarks of GVHD^14, 15^, and ITK is involved in these T cell activities. Here we used several distinct but complementary models, including *Itk_-/-_* mice, SLP76 Y145F knock-in (SLP76 Y145FKI) mice, and a newly developed ITK-specific inhibitory peptide to investigate the therapeutic potential of targeting ITK signaling in donor T cells for the separation of GVHD and GVT. Using allo-HSCT models from C57Bl/6 mice to BALB/c mice to examine GVHD and GVT by T cells from *Itk _-/-_* or SLP76 Y145FKI mice^6, 8^, we found that both CD4_+_ and CD8_+_ T cells transplanted from ITK signaling deficient mice did not induce GVHD, but retained GVT function compared to T cells from wild type (WT) mice. WT bone marrow derived T cells exhibited GVT effect but eventually caused GVHD. Furthermore, while IMP T cells derived from *Itk_-/-_* bone marrow cells were able to clear the tumor without inducing GVHD, T cells from IL-4 receptor and ITK double knockout (*Itk/Il4ra* DKO) which lack this phenotype also did not induce GVHD, indicating that absence of ITK, and not IMP cells, is responsible for reduced GVHD in the absence of ITK. *Itk_-/-_* T cells also exhibited increased expression of perforin and Eomes, which are necessary for tumor killing. In addition, T cells from *Itk_-/-_* mice showed significantly reduced expression of pro-inflammatory cytokines and decreased tissue damage during allo-HSCT. A novel and specific ITK inhibitory peptide that prevents the SH2 domain of ITK from docking onto SLP76 at tyrosine 145 position (SLP76pTYR), specifically inhibited the phosphorylation of ITK and downstream signaling molecules including PLCγ and ERK, without affecting the phosphorylation mTOR, P13K or AKT. SLP76pTYR also enhanced the development of FoxP3_+_ Treg cells. SLP76pTYR also significantly reduces IFN-γ and TNF-α by T cells from human GVHD patients. Finally, this novel specific peptide significantly reduces GVHD pathophysiology but maintained GVT function in an allo-HSCT major mismatch model. Our studies therefore identify a specific and novel potential therapeutic target for separating GVHD and GVT after allo-HSCT, with potential in other T cell mediated diseases.

## Results

### Ablation of ITK retains GVT effect but avoids GVHD during allo-HSCT

To determine whether TCR-mediated activation of ITK impacts GVHD pathogenesis after allo-HSCT, we examined the effects of ITK signaling on donor CD4_+_ and CD8_+_ T cells in allo-transplant models, utilized C57Bl/6 mice (MHC haplotype b) as donors and BALB/c mice (MHC haplotype d) as recipients. To induce GVHD, we used MHC-mismatched donors and recipient, utilizing T cell-depleted bone marrow cells from B6.PL-*Thy1_a_*/CyJ (Thy1.1) mice, along with T cells from C57BL/6 (B6) WT or *Itk_-/-_* mice injected into irradiated BALB/c mice along with B-ALL-*luc* cells. Lethally irradiated BALB/c mice were injected intravenously with 5×10_6_ wildtype (WT) T cell-depleted donor BM cells along with 1×10_6_ FACS-sorted donor T cells (1×10_6_ CD8_+_ and 1×10_6_ CD4_+_), followed by intravenous challenge with 2×10_5_ luciferase-expressing primary tumor cells B-ALL-*luc* blast cells as described_16_. Recipient BALB/c mice were monitored for tumor growth using IVIS for over 60 days **(Fig. 1A)**. While tumor growth was observed in T cell-depleted BM-transplanted mice without T cells, tumor growth was not seen in mice transplanted with T cells from either WT or *Itk_-/-_* mice. As expected, mice transplanted with WT T cells cleared the tumor but suffered significantly from GVHD. By contrast, mice transplanted with *Itk_-/-_* T cells cleared the tumor and displayed minimal signs of GVHD. Most animals transplanted with *Itk_-/-_* T cells survived for more than 65 days post-allo-HSCT **(Fig. 1B)**, with significantly reduced weight loss and clinical scores compared to those transplanted with WT T cells (scored based on weight, posture, activity, fur texture, and skin integrity as previously described^17^**, Fig. 1C-D**). BALB/c mice transplanted with WT T cells suffered from GVHD, while mice transplanted with *Itk_-/-_* T cells survived for > 65 days post-HSCT and tumor challenge with minimal signs of GVHD **(Fig. 1E)**.

**Figure 1.**
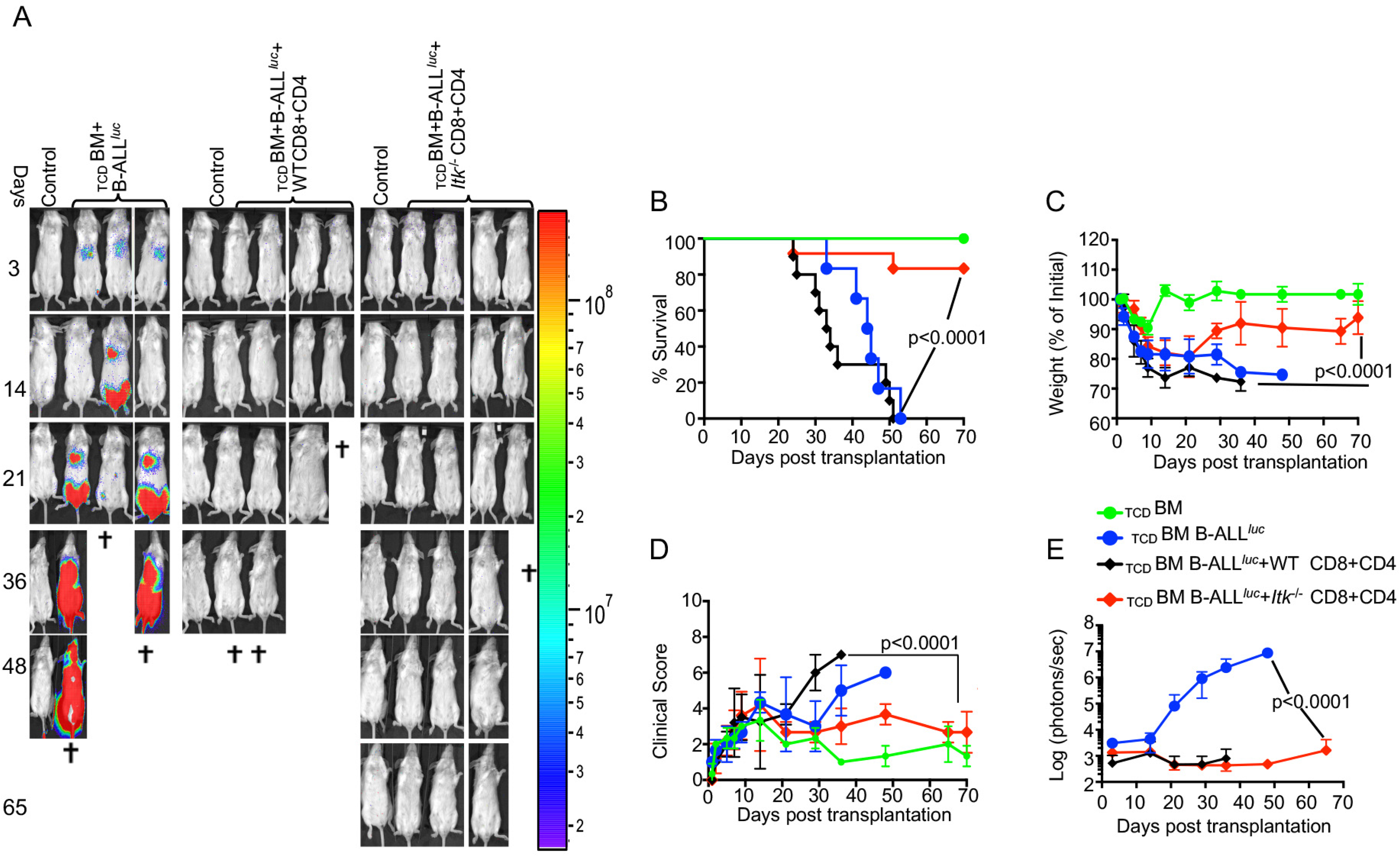
Absence of ITK retains GVT effect but avoids GVHD during allo-HSCT. 1X10_6_ purified WT or *Itk_-/-_* CD8_+_ T cells and 1X10_6_ purified CD4_+_ T cells were mixed at a 1:1 ratio, and transplanted along with 2X10_5_ B-ALL-*luc* cells into irradiated BALB/c mice. Host BALB/c mice were imaged using IVIS 3 times a week. Group 1 received T cell depleted bone marrow only (labeled as _TCD_BM). Group 2 received T cell depleted bone marrow along with 1X10_5_ B-ALL-*luc* cells (_TCD_BM+B-ALL *_luc_*+), Group 3 was transplanted with 1X10_6_ purified WT CD8_+_ and 1X10_6_ CD4_+_ T cells (1:1 ratio) along with 2X10_5_ B-ALL-*luc*+ cells (_TCD_BM+B-ALL *_luc_*+ WT CD8+CD4). Group 4 received 1X10_6_ purified *Itk_-/-_* CD8_+_ and 1X10_6_ CD4_+_ T cells (1:1 ratio) along with 2X10_5_ B-ALL-*luc*+ cells (_TCD_BM+B-ALL*_luc_*+ *Itk_-/-_* CD8+CD4). **(A)** Recipient BALB/c mice were imaged using IVIS 3 times a week. **(B)** The mice were monitored for survival, **(C)** changes in body weight, and **(D)** clinical score for 65 days post BMT. (**E**) Quantitated luciferase bioluminescence of tumor growth. Statistical analysis for survival and clinical score was performed using log-rank test and two-way ANOVA, respectively. For weight changes and clinical score, one representative of 2 independent experiments is shown (n = 3 mice/group for BM alone; n = 5 experimental mice/group for all three groups. Survival is a combination of 2 experiments. P values presented with each group. Two-way ANOVA and students t-test were used for statistical analysis*. Note: Control mouse is naïve mice used negative control for BLI*.

Donor CD8_+_ T cells are more potent than CD4_+_ T cells in mediating GVT effects, but both CD4_+_ and CD8_+_ T cells mediate severe GVHD in mice and humans^18, 19^. To determine whether CD4_+_ T cell-intrinsic ITK signaling might be sufficient to induce GVHD, we repeated the same experiments using purified CD4_+_ T cells from either WT or *Itk_-/-_* mice in the MHC-mismatch mouse model of allo-HSCT (B6 BALB/c) (**Sup. Fig. 1A-C**). Recipients of WT CD4_+_ T cells exhibited greater weight loss and mortality compared to mice receiving T cell-depleted bone marrow (_TCD_BM) cells alone (**Sup. Fig. 1B)**. In contrast, recipients of _TCD_BM mixed with *Itk_-/-_* CD4_+_ T cells had limited signs of GVHD, with greatly reduced mortality and clinical scores, indicating that CD4_+_ T cell-intrinsic ITK signaling can contribute to the severity of GVHD **(Sup. Fig. 1B-C)**. Our results indicate that ITK signaling is dispensable for anti-tumor immunity, but required for GVHD. Given the role of SLP76 in regulating ITK signaling, we next tested whether the GVT effect described above would remain intact when allo-HSCT was performed with T cells from SLP76 Y145FKI mice, which have a disrupted ITK signaling^8^. We used CD8_+_ T cells, and CD4_+_ T cells mixed at 1:1 ratio as donors as shown in Figure 1. Lethally irradiated BALB/c mice were transplanted with T cell-depleted BM (_TCD_BM) alone, or together with FACS-sorted CD8_+_ T cells from either C57Bl/6 WT or SLP76 Y145FKI mice, and challenged intravenously with B-ALL-*luc* tumor cells. While tumor growth was observed in _TCD_BM-transplanted mice without T cells, tumor growth was not seen in mice transplanted with either WT or SLP76 Y145FKI T cells. Notably, mice transplanted with WT T cells suffered from GVHD, while mice transplanted with SLP76 Y145FKI T cells survived for > 65 days post-HSCT and tumor challenge with minimal signs of GVHD and **(Sup. Fig. 2A-E)**. To determine whether tumor clearance by SLP76 Y145FKI CD8_+_ T cells was limited to B-All-*luc* cells, we performed a similar experiment using intravenously injected luciferase-expressing A20 *luc* cells^20, 21^. A20 *luc* cells failed to grow in mice that were transplanted with either WT or SLP76 Y145FKI CD8_+_ T cells (**Sup. Fig. 2).** Next, we tested purified *Itk_-/-_* or SLP76 Y145FKI CD4_+_ T cells in the same model **(Sup. Fig. 1D-F)**. When we used SLP76 Y145FKI CD4_+_ T cells, only 2 out of 10 mice developed GVHD, while all animals that received WT T cells had to be euthanized due to GVHD. Mice transplanted with SLP76 Y145FKI CD4_+_ T cells also exhibited reduced weight loss and clinical symptoms of GVHD **(Sup. Fig. 1D-F)**.

### The regulatory function of ITK in GVHD is T cell-intrinsic

The innate memory phenotype (IMP: CD44_hi_CD122_hi_) of *Itk****_-/-_*** CD8_+_ T cells arises in the thymus during development, as opposed to memory CD8_+_ T cells that are also CD44_hi_, but largely arise in the periphery of WT mice in response to foreign antigens or due to homeostatic proliferation ^13, 22^. We examined pre-transplanted CD8_+_ T cells for CD44 expression, and observed that *Itk_-/-_* CD8_+_ T cells expressed substantially higher levels of CD44 compared to CD8_+_ T cells from WT mice **(Fig. 2A).** We sought to understand whether the emergence of IMP is sufficient to separate GVHD from GVT. To test this, we generated WT IMP T cells using a mixed-bone marrow approach in which T cell-depleted BM from WT and *Itk_-/-_* mice were mixed at a 3:1 (WT: *Itk_-/-_*) ratio^23^. The irradiated syngeneic (B6) Thy1.1 hosts were reconstituted with this mixture of T cell-depleted CD45.2_+_ WT and CD45.1_+_*Itk_-/-_* BM cells, along with a control group receiving mixed CD45.2_+_ WT and CD45.1_+_ WT BM cells **(Fig. 2B)**. WT BM-derived CD8_+_ thymocytes that develop in such mixed BM chimera acquire an IMP phenotype due to their development in the same thymus as the *Itk_-/-_* T cells^23^, which we also observed in our experiments (**Fig. 2B**)^13, 22^. After reconstitution of the T cell compartment 10 weeks later, T cells derived from WT (CD45.2_+_Thy1.1_-_) and *Itk_-/-_* (CD45.1_+_) donor cells were sorted from the bone marrow chimeras.

**Figure 2.**
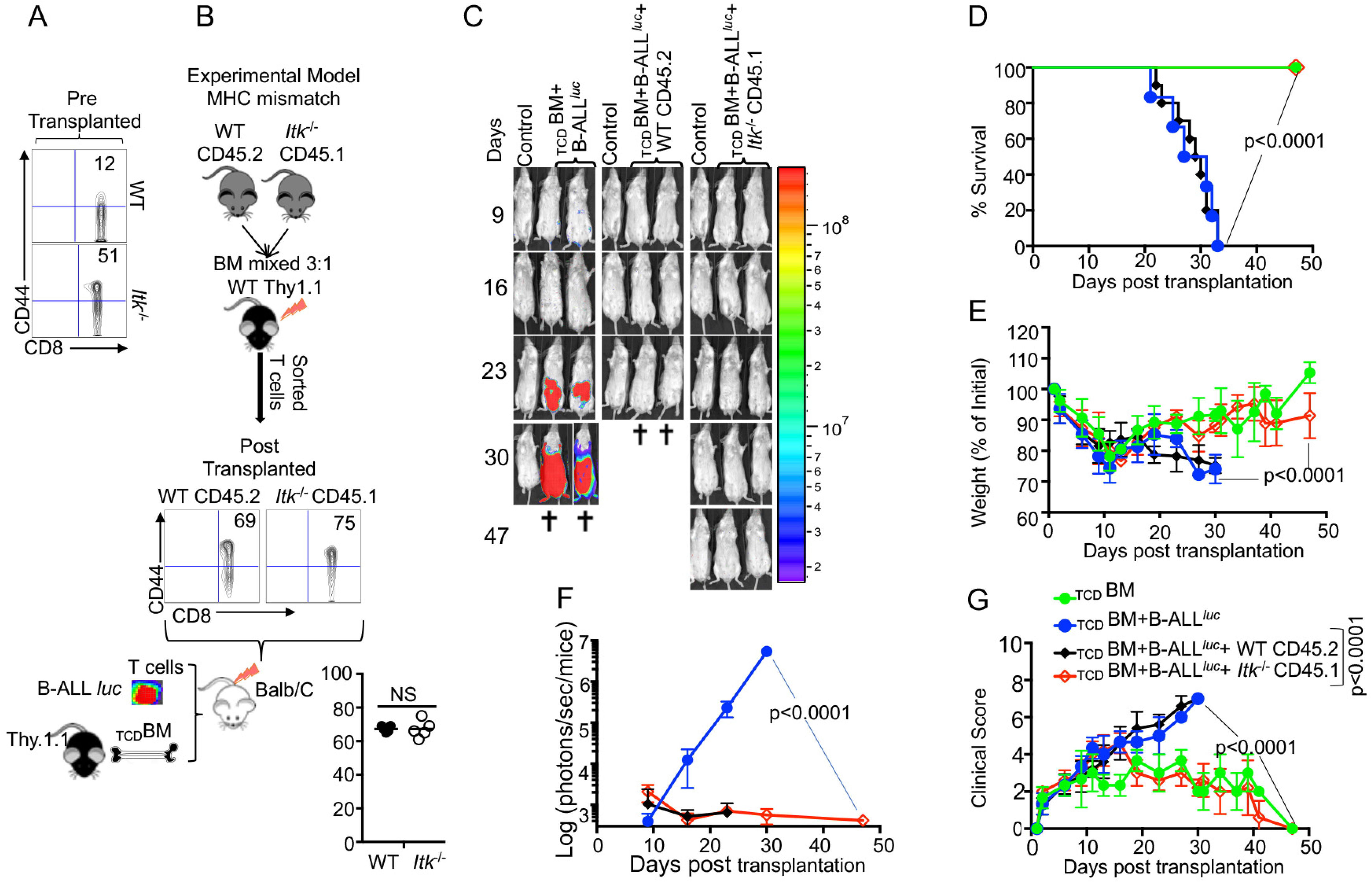
The regulatory function of ITK in GVHD is T cell-intrinsic. **(A)** Purified WT and *Itk_-/-_* CD8_+_ T cells were examined for expression of CD44 prior to transplantation. (**B**) Whole bone marrow cells isolated from C57Bl/6 WT (CD45.2) and *Itk_-/-_* (CD45.1) mice were mixed, and transplanted into irradiated Thy1.1 C57Bl/6 mice. 9-10 weeks later CD8_+_ T cells were sorted by CD45.2 and CD45.1 expression (donor T cells) and exclusion of Thy1.1 positive (host T cells). Isolated sorted T cells were examined for expression of CD44, and transplanted into irradiated BALB/c mice. This experiment were repeated three times. (**C**) Irradiated BALB/c mice were divided in four different groups and transplanted with the sorted T cells described in **(B)** as follows: Group one was transplanted with 10X10_6TCD_BM alone (_TCD_BM). Group two was transplanted with 10X10_6TCD_BM and 2X10_5_ B-ALL-*luc*, (_TCD_BM+B-ALL_luc_). Group three were transplanted with 10X10_6 TCD_BM along with 1X10_6_ purified WT 1X10_6_ CD8_+_ T cells and 2X10_5_ B-ALL-*luc* (_TCD_BM+B-ALL*_luc_*+WT CD45.2). The fourth group were transplanted 10X10_6 TCD_BM along with and 1X10_6_ purified *Itk_-/-_* CD8_+_ T cells and 2X10_5_ B-ALL-*luc* (_TCD_BM+B-ALL*_luc_*+*Itk_-/-_* CD45.1). These mice were monitored for tumor growth using IVIS system. **(D)** The mice were monitored for survival, **(E)** changes in body weight, and **(G**) clinical score for 47 days post BMT. For body weight changes and clinical score, one representative of 2 independent experiments is shown (n = 3 mice/group for BM alone; n = 5 experimental mice/group for all three groups). **(G)** Quantitated luciferase bioluminescence of luciferase expressing B-ALL-*luc* cells. Statistical analysis for survival and clinical score was performed using log-rank test and two-way ANOVA, respectively. One representative experiment out of 2. Survival is combination of 2 experiments, 3 mice per group of control T cells depleted bone marrow, and 5 mice per group for all the experimental group. P value presented with each figure. *Note: Control mouse is naïve mice used negative control for BLI*.

These sorted T cells were transplanted into irradiated BALB/c mice along with _TCD_BM in the allo-HSCT model as described above, and tested for their function in GVHD and GVT. Analysis of the BALB/c recipients of these different IMP CD8_+_ T cells indicates that WT IMP cells were not able to separate GVT from GVHD effects **(Fig. 2C-G)**. Thus, the appearance of IMP is not sufficient to separate GVHD from GVT; other mechanisms are also essential for the protective effects of ITK deficient T cells.

### Eomes expression but not innate memory phenotype T cells is required for GVT effect

*Itk_-/-_* CD8_+_ T cells exhibit attenuated TCR signaling and an innate memory phenotype (IMP)^23^, as indicated by expression of high levels of CD44, CD122, and a key transcription factor Eomes **(Fig. 3A).** To examine how these IMP T cells from *Itk*_-/-_ mice mount GVT responses, we utilized the MHC-mismatch mouse model of allo-HSCT (WT, *Itk_-/-_* BALB/c, i.e., H2K_b+_ H2K_d+_). We then sorted B6 (H2K_b+_) T cells from recipient mice and determined their cytotoxicity against B-ALL-*luc* cells. We found that these cells effectively killed primary tumor cells *in vitro*, even in the absence of ITK (**Fig. 3B**). Moreover, we observed significantly increased expression of perforin in CD8_+_ T cells from *Itk_-/-_* mice compared to T cells from WT mice in the absence of activation **(Fig. 3C)**. Our findings demonstrate that CD8_+_ T cells from *Itk_-/-_* mice have enhanced activation markers, and exert cytotoxicity against primary tumor cells.

**Figure 3.**
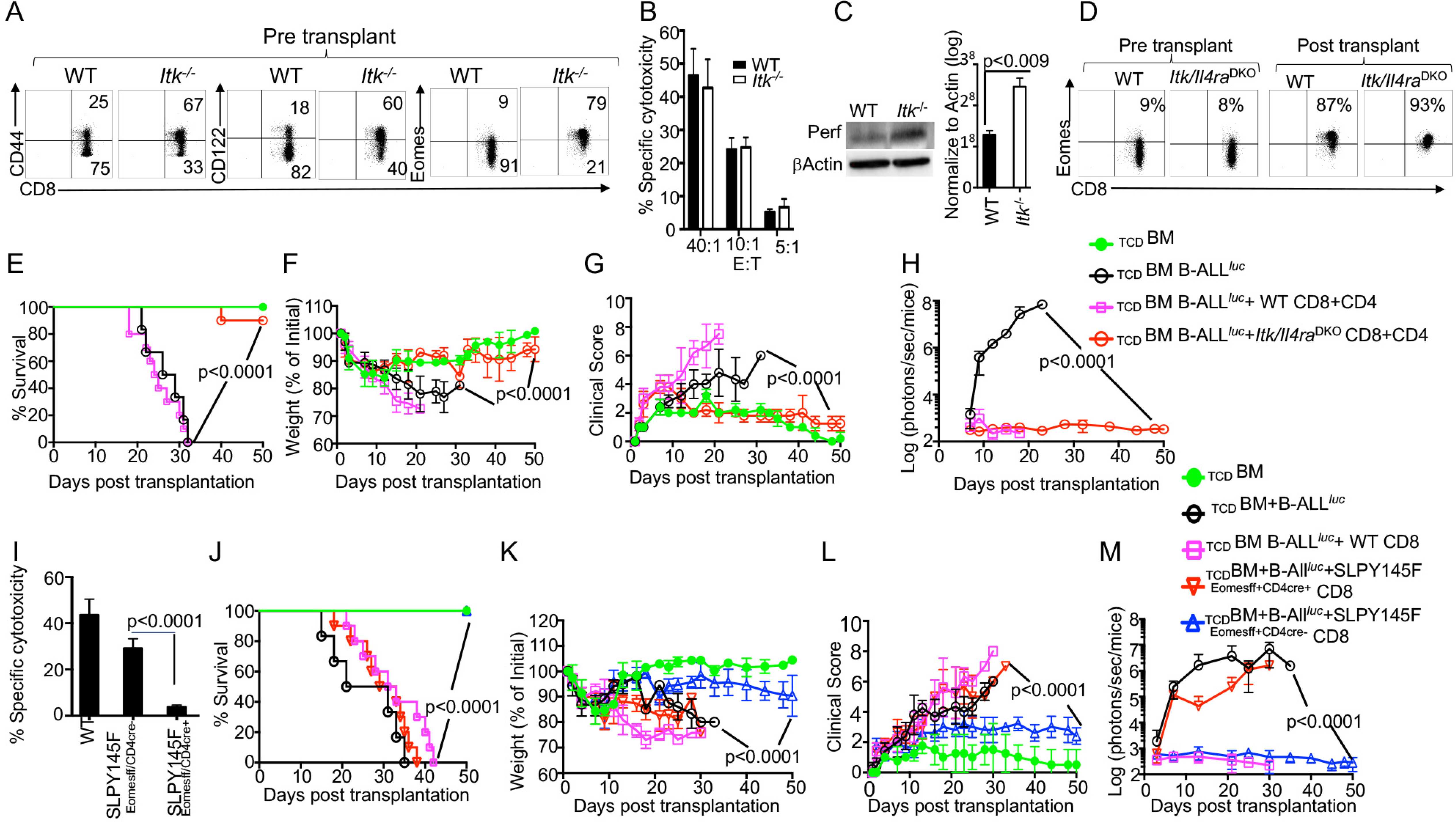
Eomes expression, but not innate memory phenotype T cells, is required for GVT effect. **(A)** Purified WT and *Itk_-/-_* T cells were examined for expression of CD44, CD122, and Eomes by flow cytometry. **(B)** Purified WT or *Itk_-/-_* T cells were transplanted into irradiated BALB/c mice, at day 7 purified T cells were sorted by H2K_d_, CD45.1 and CD45.2 expression. *Ex vivo* purified T cells were used in cytotoxicity assay against primary tumor target B-ALL*luc*+ cells at 40:1, 20:1, 5:1 ratio. **(C)** Purified T cells were examined for perforin by western blot. Quantitative analysis of four experiments of perforin expression by western blot data normalized against βActin. **(D)** *Itk/Il4ra* DKO T cells were examined for Eomes expression pre- and post-transplantation. **(E)** 1X10_6_ purified WT and *Itk/Il4ra* DKO CD8_+_ T and 1X10_6_ purified CD4_+_ T cells were mixed and transplanted along with 2X10_5_ primary B-ALL-*luc*+ cells into irradiated BALB/c mice. Host BALB/c mice were imaged using IVIS 3 times a week. Group one received T cell depleted bone marrow alone (_TCD_BM). Group two received T cell depleted bone marrow along with 1X10_5_ B-ALL-*luc* cells (_TCD_BM+B-ALL*_luc_*). Group three was transplanted with 1X10_6_ purified WT T cells and 2X10_5_ B-ALL-*luc* cells (_TCD_BM+B-ALL *_luc_ +*WT CD8+CD4). Group four received 1X10_6_ purified T cells from *Itk/Il4ra* DKO along with 2X10_5_ B-ALL-*luc* cells (_TCD_BM+B-ALL*_luc_*+ *Itk/Il4ra* DKO CD8+CD4). Recipient animals were monitored for survival, **(F)** changes in weight, and **(G)** clinical score. **(H)** Tumor burden was monitored as in figure 1. One representative of 2 independent experiments is shown (n = 3 mice/group for BM alone; n = 5 experimental mice/group for all three groups. The survival group were combinations of all experiments). **(I)** Purified donor T cells from either WT or SLP76 Y145FKI Eomes_flox/flox_ mice with or without CD4cre were transplanted into irradiated BALB/c mice. On day 7 donor T cells were purified as described and used in *ex vivo* cytotoxicity assay against B-ALL*_luc_*-cells at 40:1 ratios. **(J)** 1X10_6_ purified WT SLP76 Y145FKI Eomes_flox/flox_ with or without CD4cre CD8_+_ T cells and 1X10_6_ purified CD4_+_ T cells were mixed and transplanted along with 2X10_5_ B-ALL-*luc* cells into irradiated BALB/c mice. Host BALB/c mice were imaged using IVIS 3 times a week. Group one received T cell depleted bone marrow alone as (_TCD_BM). Group two received T cell depleted bone marrow along with 2X10_5_ B-ALL-*luc* cells (_TCD_BM+B-ALL*_luc_*). Group three were transplanted with 1X10_6_ purified WT T cells 2X10_5_ B-ALL-*luc* cells (_TCD_BM+B-ALL*_luc_+*WT CD8). Group four received 1X10_6_ purified T cells from SLP76 Y145FKI Eomes_flox/flox_ cross with CD4+cre along with 2X10_5_ B-ALL- *luc* cells (_TCD_BM+B ALL*_luc_+*SLP75Y145FKI _EomesFF+CD4cre_ CD8). Group five received 1X10_6_ purified SLP76 Y145FKI Eomes_flox/flox_ without CD4cre T cells along with 2X10_5_ B-ALL-*luc* cells (_TCD_BM+B-ALL*_luc_*+SLP75Y145FKI _EomesFF+CD4cre-_ CD8). The mice were monitored for survival, **(K)** body weight changes, and (**L)** clinical score for 50 days post BMT. For weight changes and clinical score, one representative of 2 independent experiments is shown (n = 3 mice/group for BM alone; n = 5 experimental mice/group for all three group. The survival group were a combination of all experiments. (**M**) Quantitated luciferase bioluminescence of tumor growth. Statistical analysis for survival and clinical score was performed using log-Two-way ANOVA were used for statistical analysis confirming by students *t* test, p values are presented. *Note: Control mouse is naïve mice used negative control for BLI*.

IL-4 is known to upregulate Eomes^13, 24^, which we verified by comparing T cells from WT and *Itk/Il4ra* double KO (DKO) mice. Removing IL-4 signaling from the *Itk_-/-_* mice led to decreased expression of Eomes compared to T cells from WT mice **(Fig. 3D).** Next we used the allo-HSCT model, where T cells from WT or *Itk/Il4ra* DKO were transplanted into irradiated BALB/c mice. 7 days post transplantation, WT or *Itk/Il4ra* DKO T cells were then sorted from the BALB/c recipient mice and Eomes expression by these donor T cells determined. We observed that upon allo-activation, the donor WT or *Itk/Il4ra* T cells still show increased Eomes expression **(Fig. 3D) (Sup. Fig. 3).** Next, we tested the function of *Itk/Il4ra* DKO T cells in long term allo-HSCT model. WT or *Itk/Il4ra* DKO BM were transplanted into irradiated BALB/c for GVHD and GVT studies. We observed that donor T cells from *Itk/Il4ra* DKO mice did not induce GVHD, and most of the animals survived compared to recipient mice that received donor T cells from WT mice **(Fig. 3E).** BALB/c transplanted with *Itk/Il4ra* donor T cells also had much less weight loss and significantly better clinical scores compared to BALB/c mice transplanted with WT donor T cells **(Fig. 3F-G).** Furthermore *Itk/Il4ra* DKO donor T cells cleared tumor without inducing GVHD **(Fig. 3H)**. This data demonstrated that IMP CD8_+_ cells may not be important for GVHD, but that the loss of ITK is essential for GVHD and GVT.

To further investigate the role of Eomes in tumor clearance and in the cytotoxic function we crossed SLP76Y145FKI mice with Eomes_flox/flox_ and crossed these mice with CD4cre to delete the Eomes specifically in T cells^10, 25^. To obtain ex *vivo* activated cells, we performed similar allo-HSCT experiments as described above, and additionally used T cells from WT or SLP76Y145FKI mice with or without Eomes. 7 days post-transplant, donor T cells were sorted as described by H2K_b_ positivity, and *in vitro* cytotoxicity assays were performed at 40:1 ratios. We observed that donor T cells lacking Eomes were unable to kill tumor targets **(Fig. 3I).** Next, we examined the role of Eomes in the allo-HSCT model. Lethally irradiated BALB/c mice were injected intravenously with 5×10_6_ WT T cell-depleted BM cells along with 1×10_6_ FACS-sorted CD8_+_ and CD4_+_ from either WT mice or SLP76Y145FKI Eomes_flox/flox_ with or without CD4cre, followed by intravenous challenge with 2×10_5_ luciferase-expressing primary tumor cells B-ALL-*luc* blast cells as described ^16^. Recipient animals transplanted with WT T cells cleared the tumors cells but developed acute GVHD **(Fig. 3J).** Recipient animals transplanted with T cells from SLP76Y145FKI Eomes_flox/flox_ without CD4cre mice, however, cleared the tumors without showing signs of GVHD **(Fig. 3L)** and **(Sup. Fig. 3B)..** Notably, recipient animals transplanted with T cells from SLP76Y145FKI Eomes_flox/flox_ with CD4cre mice were unable to clear the tumor and all died from tumor burden. This data provided further evidence that Eomes is required for the GVT effects **(Fig. 3M).**

### ITK deficiency results in reduced cytokine production

It is known that the conditioning regimen for allo-HSCT elicits an increase in the production of inflammatory cytokines by donor T cells, known as a “cytokine storm”, and is considered one of the hallmarks of GVHD pathogenesis^26^. We obtained blood samples from GVHD patients and examined the levels of serum inflammatory cytokines such as IL-33, IL-1α, IFN-γ, TNF-α and IL-17A compared to healthy donors. We observed that patients with GVHD have significantly higher level of serum proinflammatory cytokines compared to healthy control (**Fig. 4A**). Next we assessed cytokine production by *Itk_-/-_* T cells in our allo-HSCT model (B6 BALB/c), examining the levels of serum inflammatory cytokines such as IL-33, IL-1α, IFN-γ, TNF-α and IL-17A on day 7 post allo-transplantation **(Fig. 4B).** We found that serum IFN-γ and TNF-α were significantly reduced in recipients that received *Itk_-/-_* CD8_+_ T cells mice compared to those that received WT CD8_+_ T cells **(Fig. 4B)**. Thus, we confirmed that the findings in our pre-clinical model correlated with human GVHD samples. We also isolated *Itk_-/-_* donor T cells from the secondary lymphoid organs of recipients using anti-H2K_b_ antibodies (expressed by donor C57Bl/6 cells) and stimulated them with anti-CD3/CD28 **(Fig. 4C),** or PMA/ionomycin (to bypass the proximal signaling defect, (**Sup. Fig. 4**) in the presence of Brefeldin A, or left them unstimulated for 6 hours, followed by analysis of IFN-γ and TNF-α cytokine production. Compared to WT T cells, *Itk_-/-_* T cells were capable of producing IFN-γ and TNF-α when T cell signaling was bypassed by re-stimulation with PMA and ionomycin (**Sup. Fig. 4**), however, they produced significantly less inflammatory cytokines when stimulated via TCR/CD28 **(Fig. 4C, D).** Next, we determined whether the reduction of cytokine production by *Itk_-/-_* donor T cells was due to cell-intrinsic or extrinsic factors. We mixed purified *Itk_-/-_* CD8_+_ T cells with purified WT CD8_+_ T cells at a 1:1 ratio and transplanted the mixed cells into irradiated BALB/c as described above. On day 7, donor T cells were isolated from recipient mice using H2K_b+_ and examined for IFN-γ and TNF-α expression as described above. We found that WT donor CD8_+_ T cells produced higher levels of inflammatory cytokines than *Itk_-/-_* donor CD8_+_ T cells, suggesting that the reduced cytokine production observed by *Itk_-/-_* donor T cells is T cell-intrinsic **(Fig. 4E**).

**Figure 4.**
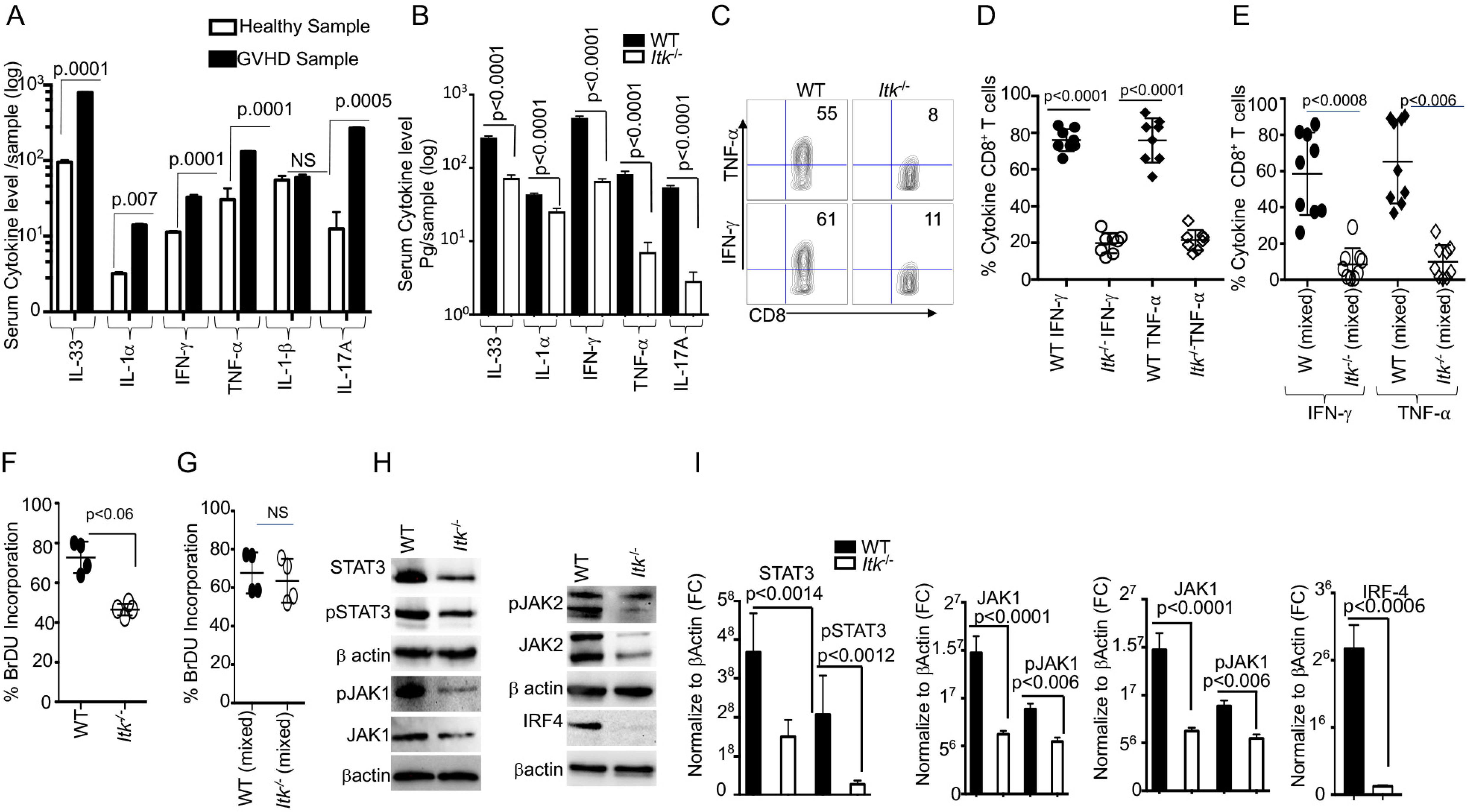
ITK deficiency results in reduced cytokine production. **(A)** Serum from Several GVHD patients were Isolated and examine for inflammatory cytokines production (IL-33, IL1α, IFN-γ and TNF-α and IL-17A) determined by ELISA (**B**) 1X10_6_ purified WT or *Itk_-/-_* CD8_+_ T cells were transplanted with into irradiated BALB/c mice. **(B)** At day 7 the post allo-HSCT, recipient BALB/c were euthanized and serum cytokines (IL-33, IL1α, IFN-γ and TNF-α and IL-17A) determined by ELISA. **(C, D)** Intracellular IFN-γ and TNF-α expression by donor CD8_+_ T cells after stimulation with anti-CD3/anti-CD28 as determined by flow cytometry. Combined data from 3 independent experiments is shown for (C). **(E)** Flow cytometry analysis of purified WT and *Itk_-/-_* T cells that were mixed at a 1:1 ratio for transplantation into irradiated BALB/c mice. At day 7 donor T cells were gated for expression of H-2K_b_, CD45.1, and CD45.2 and analyzed intracellular expression of IFN-γ and TNF-α by flow cytometry after stimulation with anti-CD3/anti-CD28. Combined data from 4 independent experiments is shown, and the p values for each experiment is shown. (**F**) Purified WT or *Itk_-/-_* donor T cells were transplanted into irradiated BALB/c. At day 7 donor cells were analyzed for donor T cells proliferation by examining BrDU incorporation by flow cytometry. **(G)** Purified WT and *Itk_-/-_* donor T cells were mixed 1:1 ratio and transplanted into irradiated BALB/c, at day 7 donor T cells were gated for the expression of H-2K_b_, CD45.1, and CD45.2 and analyzed for BrDU incorporation. **(H)** Purified WT and *Itk_-/-_*T cells were examined for the expression and phosphorylation of IRF4, JAK1/2 and STAT3 by western blot. **(I)** Quantitative analysis from western blots using ImageJ to normalize to βActin, data from 3 independent experiments. For statistical analysis we used two-way ANOVA and student’s *t* test, p values are presented.

We next examined donor T cell proliferation using a BrdU incorporation assay. 7 days post allo-transplantation as described above, transplanted T cells were examined for proliferation by BrdU incorporation. Donor T cells from *Itk_-/-_* mice showed reduced proliferation compared to donor T cells from WT mice **(Fig. 4F).** To determine if the reduced proliferation of *Itk_-/-_* donor T cells was due to cell-intrinsic mechanisms, we mixed sort purified *Itk_-/-_* CD8_+_ T cells with purified WT CD8_+_ T cells at a 1:1 ratio, followed by transplantation as described above. Interestingly, no difference was observed in BrdU incorporation between WT and *Itk*_-/-_ donor T cells in the mixed transplant models, indicating that the reduced proliferation of donor *Itk_-/-_* T cells proliferation was due to cell-extrinsic effects **(Fig. 4G)**. Thus, both cell intrinsic and extrinsic mechanisms regulate the behavior of *Itk*_-/-_ CD8_+_ donor T cells.

The transcription factor IRF4 has been shown to play critical roles in modulating TCR signaling, including TCR signal strength such as those regulated by ITK^27, 28^. The JAK/STAT signaling pathway is also critical for the response of T cells to cytokines^29, 30, 31^. To examine whether there was a differential signaling between WT and *Itk_-/-_* donor T cells in the GVHD and GVT model as determined by expression of IRF4 and JAK/STAT, we examined expression of IRF4, JAK1, JAK2 and STAT3 by purified T cells from spleen. Our data showed that *Itk_-/-_* donor T cells expressed significantly less IRF4, JAK1, JAK2, and STAT3 as well as phosphorylated forms of JAK1, JAK2 and STAT3 **(Fig. 4H, I).** Our data suggest that the lack of ITK expression affects the expression of IRF4, and the amount of cytokine signals the cells received. These data may explain the reduced cytokine production and proliferation in *Itk_-/-_* T cells observed above.

### ITK/SPL76-Y145 signaling axis is required for T cell migration to the GVHD target tissues

GVHD involves early migration of alloreactive T cells into the target organs, followed by T cell expansion and tissue destruction. Modulation of alloreactive T cell trafficking has been suggested to play a significant role in ameliorating experimental GVHD^32^. Therefore, we examined the trafficking of donor T cells to GVHD target tissues as previously described^32^. Irradiated BALB/c recipient mice were injected with CD8_+_ T cells from *Itk_-/-_* (CD45.2_+_) and WT C57Bl/6 (CD45.1_+_) mice mixed at a 1:1 ratio **(Fig. 5A)**, and 7 days post transplantation, recipient mice were examined for the presence of donor T cells in the spleen, lymph nodes, liver and the small intestines. While the WT: *Itk_-/-_* T cell ratio remained ∼1:1 in the spleen and lymph nodes **(Fig. 5B)**, this ratio in the liver and small intestine was significantly elevated suggesting that *Itk_-/-_* T cells were defective in migration to and/or expansion in those tissues. Using histological staining for H&E, we also observed significant leukocyte infiltration into GVHD target organs, liver skin and small intestine (SI)^33^, in WT T cell recipients but not in *Itk_-/-_* T cell recipients **(Fig. 5C)**. As an alternative approach, we tracked T cells in allo-BMT mice by using donor CD8_+_ T cells from WT and *Itk_-/-_* mice that also express luciferase, which could be monitored by bioluminescence ^27^. We observed that donor T cells from *Itk_-/-_* had significantly impaired residency in GVHD target organs, including the liver and small intestine (SI), compared to WT, despite no differences in spleen and lymph nodes **(Fig. 5D)**^33^. We next examined donor T cell proliferation as described above. 7 days post allo-transplantation as described above, transplanted donor T cells in liver and small intestine of recipients were examined for proliferation by BrdU incorporation. Donor T cells from *Itk_-/-_* mice showed reduced proliferation compared to donor T cells in GVHD target organs from WT mice **(Fig. 5E).** In the mixed T cell transfer model, we had determined that *Itk_-/-_* T cell proliferation was comparable to that of WT cells; therefore, it is very likely that the reduced numbers of *Itk_-/-_* T cells in the liver and small intestine was due to impaired T cell trafficking. Pro-inflammatory conditioning treatment may promote T cell migration into GVHD target tissues ^34, 35^. Indeed, in the same mixed T cell transfer model, we found that chemokine and chemokine receptor expression (Aplnr, Cxcr5, Accr2, CCL12, CCL2, CCL5, Ccr9, Ackr4, and Cmtm4) was also significantly reduced in *Itk_-/-_* CD8_+_ T cells at day 7 post-transfer **(Fig. 5F).** These data suggest that *Itk_-/_*_-_ CD8_+_ T cells display attenuated chemokine receptor expression, which correlates with defective migration to GVHD target organs and target organ pathology.

**Figure 5.**
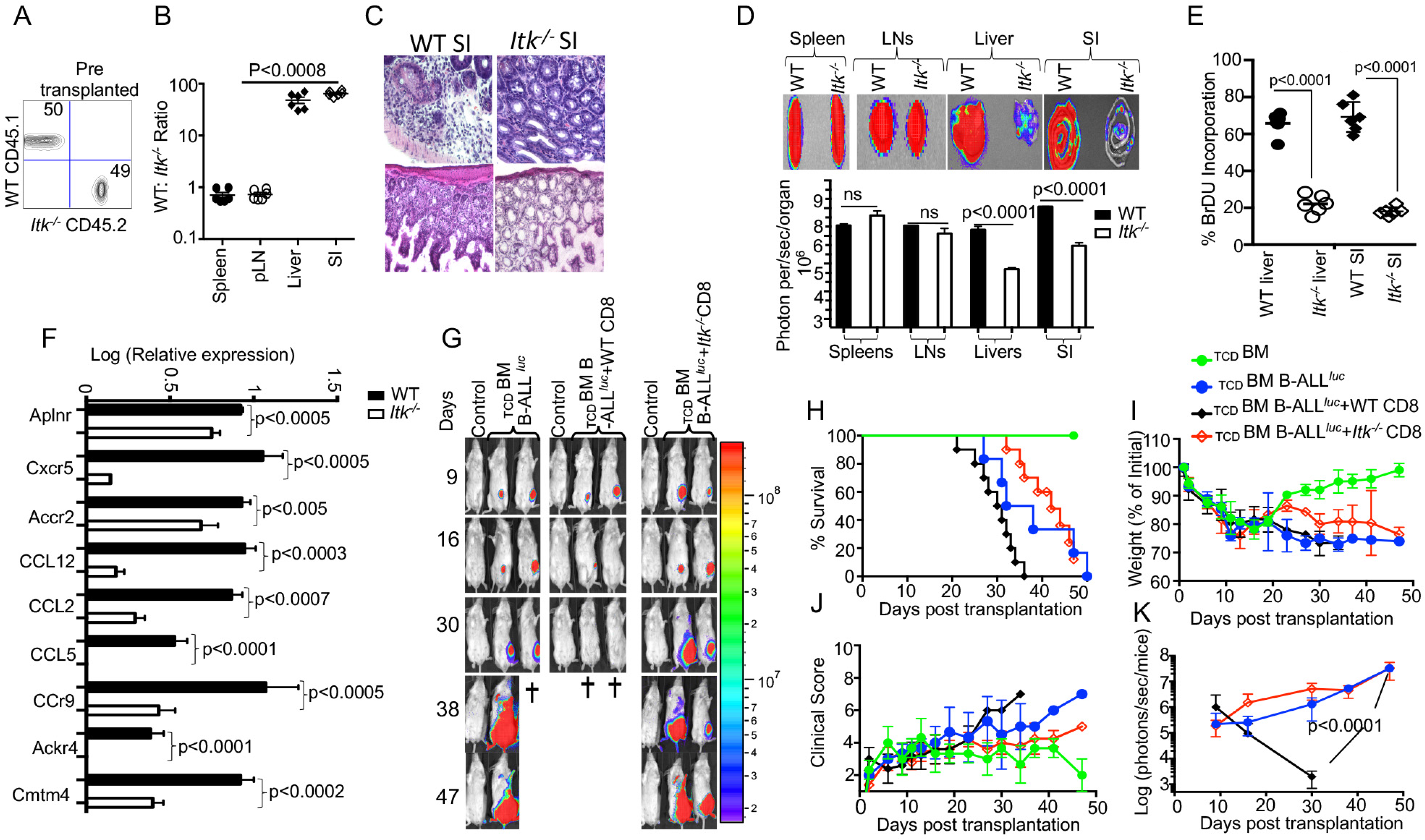
ITK/SLP76-Y145 signaling axis is required for T cell migration to the GVHD target tissues. **(A**) Irradiated BALB/c mice were allo-HSCT-transplanted and injected with FACS-sorted WT and *Itk_-/-_* CD8_+_ T cells mixed at a 1:1 ratio. FACS analysis of sorted T cells pre-transplant shown. **(B)** At day 7 post-BMT, the spleen, liver, and small intestine (SI) were examined for donor WT and *Itk_-/-_* T cells. The ratio of WT: *Itk_-/-_* CD8_+_ T cells in the organs was determined. **(C)** At day 7 post-allo-HSCT, small intestines were examined by H&E staining. **(D)** Irradiated BALB/c mice were BM-transplanted and injected with CD8_+_ T cells from luciferase-expressing WT or *Itk_-/-_* mice. Total bioluminescence from the spleen, pLN, liver, and SI, and representative bioluminescence images are shown on day 7 post-HSCT, and quantified bioluminescence. **(E)** Purified WT or *Itk_-/-_* donor T cells were transplanted into irradiated BALB/c. At day 7 donor cells were analyzed for donor T cell proliferation in recipient liver and small intestine (SI) by examining BrDU incorporation by flow cytometry. **(F)** On Day 7 post-allo-HSCT, donor CD8_+_ T cells were isolated and examined for the expression of Aplnr, Cxcr5, Accr2, CCL12, CCL2, CCL5, CCr9, Ackr4, and Cmtm4 using q-RTPCR. p values were calculated using 2-way ANOVA and Student’s *t* test, p values are listed. **(G**) Irradiated BALB/c mice were transplanted with C57Bl/6- derived BM and FACS-sorted WT or *Itk_-/-_* CD8_+_ T cells, and challenged subcutaneously with luciferase-expressing B-All *luc* cells. Recipient animals were monitored for weight changes. Group one of recipient mice were transplanted T cells depleted bone marrow (_TCD_ BM). The second group of recipient mice were transplanted T cells depleted bone marrow and primary B-ALL luciferase (_TCD_BM+B-ALL*_luc_*). The third group of recipient mice were transplanted WT T cells along 2X10_5_ primary tumor cells B-ALL-*luc* (_TCD_BM+B-ALL*_luc_*+WT CD8). The fourth group of recipient mice were transplanted with T cell depleted bone marrow and primary B-ALL-*luc*, along with *Itk*_-/-_ T cells (_TCD_BM+B-ALL*_luc_*+ *Itk*_-/-_ CD8). Representative bioluminescence images of tumor-bearing mice on days 9, 16, 30, 38, and 47 are shown. *Note: Control mouse is naïve mice used negative control for BLI.* **(H)** Animals were monitored for survival over 47 days. **(I)** Changes in weight loss. **(J)** Animals were monitored for clinical score. **(K)** Recipient mice were monitored for tumor growth using the IVIS imaging system and quantified. For weight changes and tumor burden, one representative of 2 independent experiments is shown (n = 3 mice/group) for control and n=5 mice for WT and n=5 mice for *Itk_-/-_.* Survival group was combined from both experiments. P values were calculated using two-way ANOVA and Student’s *t* test, p values are listed.

Given that *Itk_-/-_* T cells exhibit defective migration to target organs of GVHD, we predicted that although *Itk_-/-_* T cells can clear tumor cells in the blood and secondary lymphoid organs, they would not be able to kill tumors that reside in tissues. To test this possibility, lethally irradiated BALB/c mice were BM-transplanted together with FACS-sorted WT or *Itk_-/-_* CD8_+_ T cells, and challenged with subcutaneously injected B-All *luc* cells. Although *Itk_-/-_* CD8_+_ T cells did not cause GVHD, the subcutaneously injected tumors were cleared in mice transplanted with WT CD8_+_ T cells but not with *Itk_-/-_* CD8_+_ T cells **(Fig. 5G-K).** Together, these data suggest that the ITK signaling in T cells can separate GVHD from GVT effects, but only for tumors that reside in the circulation and in secondary lymphoid organs (such as hematologic malignancies).

### ITK differentially regulates gene expression in T cells during GVHD

As an unbiased approach to further explore differences between WT and *Itk_-/-_* CD8_+_ T cells, we employed RNA sequencing analysis to examine the differences in gene expression between WT and *Itk_-/-_* CD8_+_ T cells following allo-HSCT. We sort purified donor WT and *Itk_-/-_* CD8_+_ T cells (using H-2K_b_ antigen expressed by donor T cells) before and 7 days after they were transferred into irradiated BALB/c recipients, for RNA sequencing. Although WT and *Itk_-/-_* CD8_+_ T cells are distinct prior to transplantation due to the enhanced IMP in the absence of ITK, WT and *Itk_-/-_* cells homed to the spleen post transplantation are similar as revealed by the fact that they clustered within a close proximity in the Principal Component Analyses (PCA) (**Fig. 6A**). We were unable to collect enough cells from the intestine of the *Itk_-/-_* T cell recipients, since they are deficient in homing to the intestine (**Fig. 5E**), however the WT T cells that home to the intestine (referred to as “Gut”) exhibited significantly different transcriptomic profiles compared to *Itk_-/-_* as revealed by the PCA (**Fig. 6A**). To further determine the differentially expressed genes that are unique in WT cells associated with their ability to home to the GVHD target organ (intestine or “Gut”), we compared the lists of genes that were up- or down-regulated after the cells were transferred into the recipients and homed to different organs. We found that genes that are up- or down-regulated in *Itk_-/-_* T cells isolated from the spleens of the recipients (*Itk_-/-_*-Spl), as compared to *Itk_-/-_* pre-transplanted cells (*Itk_-/-_*-Pre), have minimal overlap with those that are differentially expressed in WT cells homed to the gut (normalized to WT-Pre) (**Fig. 6B&C**). Genes that are differentially expressed in WT T cells that were able to home to the GVHD target organ may reveal signals that are deficient due to the absence of ITK. We therefore extracted the list of genes that are up- or down-regulated in only WT T cells isolated from the gut of the recipients post transplantation (**Fig. 6D** shows 23 up-regulated and **Fig. 6E** shows 27 down-regulated genes). The differentially expressed genes between WT and *Itk*_-/-_ donor T cells were enriched for transcripts encoding lymphocyte homing molecules such as adhesion molecules and chemokine signaling proteins, which might contribute to the defective homing capability of *Itk*_-/-_ donor T cells (**Fig. 6F**). The results of critical gene that were differentially expressed were confirmed by q-RT-PCR (**Fig. 6G**). Using pathway enrichment analyses, our data also revealed a critical role for ITK in regulating genes involved in T cell cytokine/cytokine receptor interaction, cell adhesion, graft-versus-host disease, allograft rejection, and chemokine signaling pathways (**Fig. 5F**). These data suggest that ITK regulates the expression of signature genes associated with the homing of the transplanted cells into the GVHD targeted organs, while it does not have an apparent effect in T cell homing in the spleen. This may, in part, explain the ability of *Itk_-/-_* T cells to maintain GVT effects while being unable to home to the GVHD target organs and participate in GVHD.

**Figure 6.**
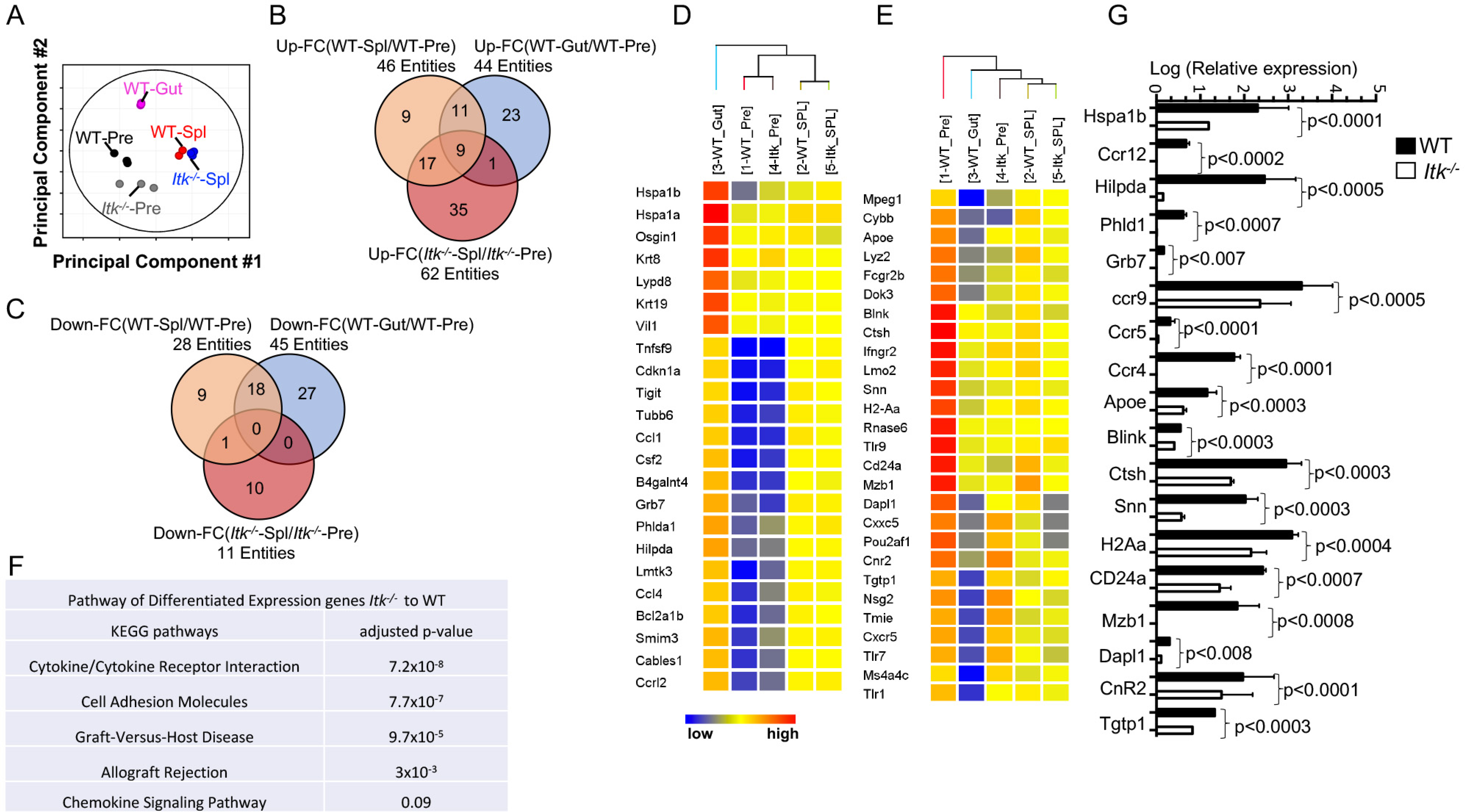
ITK differentially regulates gene expression in T cells during GVHD. WT and *Itk_-/-_* CD8_+_ T cells were FACS sorted then transplanted into irradiated BALB/c mice. At day 7 post-transplant, donor T cells were sort isolated (based on FACS expression of H-2K_b_, CD3 and CD8) from host spleen and small intestine (Gut). Sorted donor T cells were subjected to RNA sequencing. **(A)** Principal component analysis of genes with ≥ 2-fold change in any pairs of group combinations, with false discovery rate (FDR) ≤ 0.05. WT-Pre and *Itk_-/-_*-Pre-denote cells prior to transfer, and WT-Spl, WT-Gut, and *Itk_-/-_*-Spl denote cells isolated from the spleen (Spl) or small intestine (Gut) of the recipients post transfer. (**B&C**) Venn diagram of genes that are ≥ 2-fold up- or down-regulated in the indicated comparisons, with FDR (*P*) ≤ 0.05. (**D-E**) Heat map of 23 genes that are up-regulated in WT-Gut only. (**F**) Differentially expressed genes were enriched for pathway analysis comparing WT and *Itk_-/-_.* (**G**) WT and *Itk_-/-_* CD8_+_ T cells were FACS sorted then transplanted into irradiated BALB/c mice. At day 7 post-transplant, donor T cells were sort isolated (based FACS expression of H-2K_b_, CD3 and CD8) from host spleen and small intestine (Gut). Total RNA was isolated from sorted donor T cells were and qPCR performed.

#### Novel peptide SLP76pTYR inhibitor specifically targets ITK signaling and enhances Treg cell development

Since *Itk_-/-_* T cells can separate GVHD from GVT, as can SLP76 Y145FKI T cells, we sought to disrupt ITK signaling with pharmacological agents. When we used several commercially available small molecule inhibitors, including 10n ^36, 37^ and commercially available CTA056 ^38^ and GSK2250665A^39^, we observed that these small molecules also inhibit several other kinases including mTOR and AKT, suggesting that these molecules were not specific **(Sup. Fig. 5).** We thus sought to design a novel inhibitor that specifically targets ITK signaling by preventing the SH2 domain of ITK from docking onto SLP76 at tyrosine 145 position. In our quest to generate a potent inhibitor of the ITK-SH2 domain:SLP76-pY145 interaction, we analyzed the physico-chemical and structural properties of the interface of the SH2 domain and SLP76-pY145 (**Fig. 7A,B**), since evolution usually selects for residues at specific protein: protein interfaces for certain properties^40^. To design a short peptide that can inhibit ITK via competitive bidding, we examined SLP76-Y145 region of SLP76 in our initial peptide inhibitor design. In addition, to avoid the unintended non-specific binding of our peptide to the more than hundred other SH2 domains (and/or to other unexpected targets) in vivo^41^. we incorporated as many distinctive features of the SLP76 region around the pY145 as possible, using as a guide, atomic resolution NMR spectroscopy structures of the SH2 domain of ITK, free and in complex with a short peptide containing a pTyr residue^40^ **(Fig. 7A**). The SH2 domain of ITK contains complementary electrostatic surface, because the phosphotyrosine binding pocket as well as the surrounding surface groove are highly positively charged, suggesting that electrostatics most likely will play a key role in this interaction. We thus designed a novel SLP76145pTYR peptide to bind to the ITK SH2 domain in order to prevent ITK from docking onto SLP76 at tyrosine at 145 position. BLASTing^42^ the peptide sequence of our novel peptide SLP76145pTYR peptide against the non-redundant human proteome also shows minimal identity with other proteins, suggesting that SLP76145pTYR and the SH2 domain interaction is unique, and most likely will be specific towards ITK signalling. To test this, we cultured T cells with FITC-conjugated SLP76pTYR, (FITC Dye) -132NEEEEAPVEDDAD**pY**EPPPSNDEEA155-(GRKKRRQRRRPQ) vehicle or nonspecific peptide (FITC Dye) IIMTTTTNNKKSSRRVVVVAAAADD (GRKKRRQRRRPQ) and examined for FITC uptake using microscopy and flow cytometry. We observed that significant numbers of cells were positive for FITC **(Fig. 7B, D-G).** Next we examined whether the FITC label was localized in specific locations in the cell. We imaged the cells in a single focal plane near the cover glass, and observed that FITC was clustered in specific locations of the cell **(Fig. 7E)**. ITK deficiency is known to enhance the development of regulatory T cells^43, 44^, and we tested the peptide inhibitor to determine whether inhibition of ITK with SLP76pTYR would induce Tregs. Total mouse T cells were stimulated with anti-CD3^45^ and in the presence of either SLP76pTYR or nonspecific peptide for 24 to 48 hours, and cells were harvested and examined for the presence of Tregs (CD4_+_CD25_+_FoxP3_+_). We observed significantly enhanced differentiation of Treg cells in T cell cultures treated with SLP76pTYR peptide compared to vehicle alone or nonspecific peptide **(Fig. 8A-B).** Next, we examined the signaling molecules that are specifically activated upon activation of ITK, including the phosphorylation of ITK, PLCγ1, and ERK, and we observed significant reduction in the phosphorylation of these molecules compared to either vehicle alone or scramble peptide **(Fig. 8C-D)**. Next we investigated the effects of SLP76pTYR peptide on human PBMC samples from GVHD patients and normal donors. T cells from these patients were stimulated with anti-CD3 (OKT3) for 24 hours in the presence of SLP76pTYR or nonspecific peptide. We observed there was significant reduction in the phosphorylation of PLCγ1, slight reduction in pERK but no effect on pAKT and pMTOR (**Fig. 8E-F**), providing evidence that SLP76pTYR peptide has effects on both mouse and human T cells. Notably, SLP76pTYR peptide exhibited minimal off-target effects against other kinases including the mTOR, PI3K, and AKT (**Fig. 8C-F**). Next we investigated the effects of SLP76pTYR peptide on the ability of human PBMC to produce proinflammatory cytokines. T cells from normal donors were stimulated with anti-CD3 (OKT3) and CD28 in the presence of SLP76pTYR or vehicle alone. We also examined T cells incubated with SLP76pTYR or vehicle alone in the presence of PMA+I in the presence of Brefeldin A. Our data shows that T cells stmulated with anti-CD3/CD28 show significantly reduced IFN-γ and TNF-α when incubated in the presence of SLP76pTYR compared to those in the presence of vehicle alone (**Fig. 8G**). The cells also did not exhibit signs of apoptosis after 48 hours incubation with SLP76pTYR peptide, suggesting that this peptide does not induce general toxicity in the cells **(Sup. Fig. 6)**.

**Figure 7.**
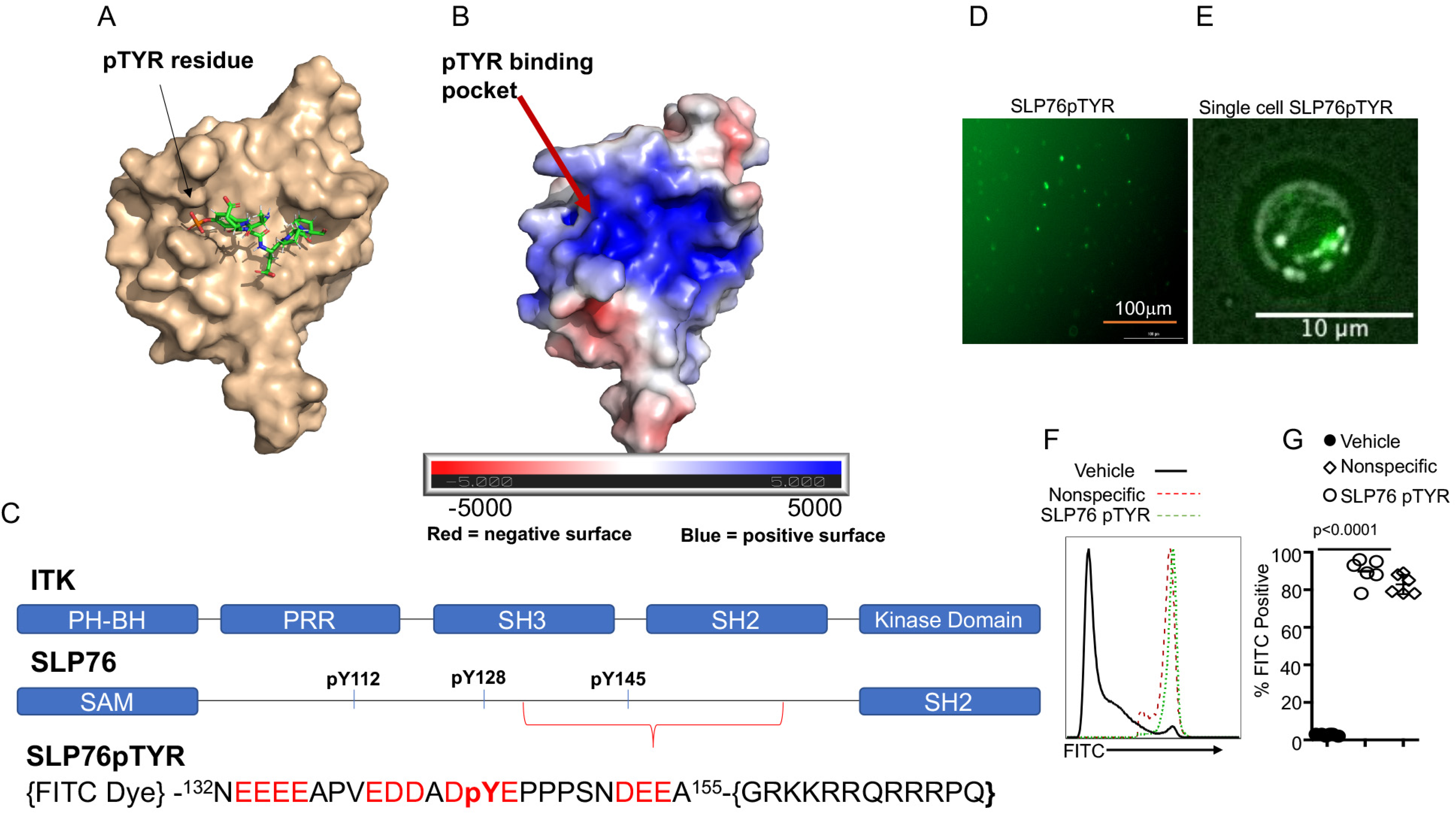
Development of a peptide that disrupts the interaction between SLP76 and ITK. **(A)** NMR Spectroscopy structures of murine ITK SH2 domain showing its complex with SLP76pTYR peptide containing a pTyr residue. The SH2 domain is rendered in surface representation (wheat), while the peptide with sequence (ADpYEPP) is shown in stick model (PDB code:2ETZ^40^), and **(B)** Electrostatic profile shown, calculated using the APBS plugin in Pymol. **(C**) Top: Organization of the domain architecture of ITK, and amino acid 132-152 of SLP76 adapter protein respectively. Bottom: Design of the novel peptide, SLP76pTYR, that inhibits the SLP76-mediated ITK signaling. **(D)**. T cells were examined for percentage FITC positive by fluorescence microscopy. **(E)** A single cell in focal plane near the cover glass, was imaged. **(F)** Primary cells cultured with SLP76pTYR or scrambled peptide were washed and examined for FITC expression by flow cytometry. **(G)** Quantification of the expression in **(F).** For statistical analysis two-way ANOVA and confirmation by Student’s *t* test were performed.

**Figure 8.**
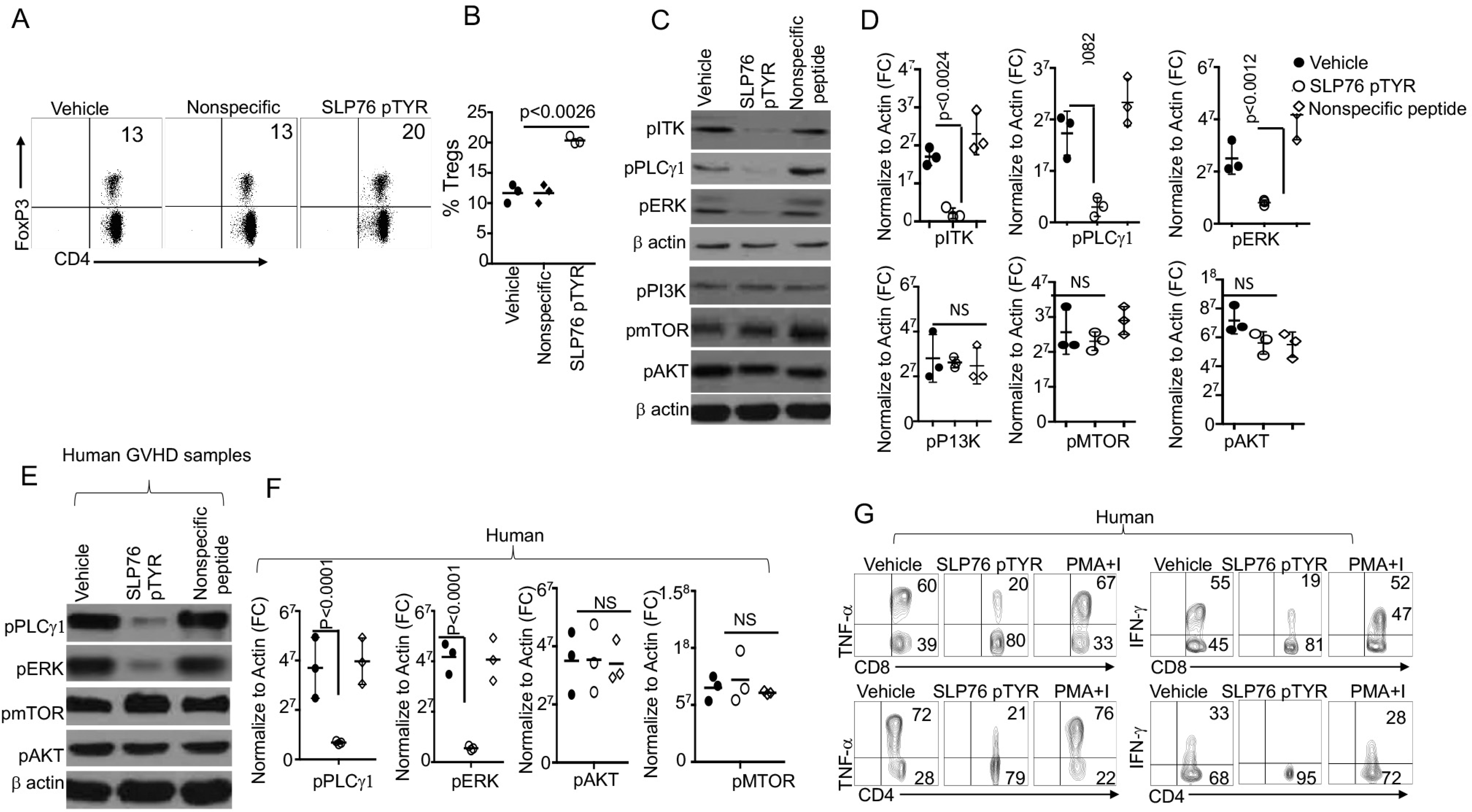
Novel peptide SLP76pTYR specifically targets ITK signaling and enhances Treg cell development. **(A)** Total T cells stimulated in the presence of SLP76pTYR or vehicle alone were examined for CD4_+_ and FoxP3_+_. n=3 one representative experiment shown. **(B)** Quantification of three experiments as in (A). **(C)** Cell lysates from T cells stimulated in the presence of SLP76pTYR or vehicle alone were examined for phosphorylation of ITK, PLCγ1 and ERK. Lysates were also examined for phosphorylation of PI3K, mTOR, P13K, and AKT. n=3 one representative experiment shown. (**D**) Western blots were normalized to β−Actin and quantitative data from 3 independent experiments presented. (**E**) Human GVHD patients’ samples were cells stimulated in the presence of SLP76pTYR or vehicle only. T cell lysates were used in western blots for analysis of pPLCγ, pERK, pAKT, pMTOR, and **(H)** three experiments were quantitated and normalized to β−Actin. n=3, one representative experiment shown. (**G**) Primary human T cells purified from PBMC were stimulated with CD3 and CD28 for 5 hours in the presence of vehicle alone or SLP76pTYR in the presence BFA. Intracellular IFN-γ and TNF-α expression by CD8_+_ and CD4_+_ T cells was determined by flow cytometry. For statistical analysis we used two-way ANOVA and Student’s *t* test. P values are presented.

#### Inhibition of T cells by peptide SLP76pTYR allows tumor clearance without inducing GVHD

Next we evaluated the potential for peptide inhibitor of SLP76pTYR in modulating Itk in vivo n GVT vs GVHD as proof of principle for the approach. WT T cells CD8_+_ T and CD4_+_ T cells were mixed at 1:1 ratio transduced with retrovirus carrying SLP76pTYR or empty vector. Lethally irradiated BALB/c mice were transplanted with T cell-depleted BM (_TCD_BM) alone, or together with SLP76pTYR (or vector) transduced WT CD8_+_ and CD4_+_ T cells, and challenged intravenously with B-ALL-*luc* tumor cells. While tumor growth was observed in _TCD_BM-transplanted mice without T cells, tumor growth was not seen in mice transplanted with either untransduced T cells or T cells transduced with either empty viruses or SLP76pTYR. Notably, mice transplanted with untransduced T cells or T cells transduced with empty viruses suffered from GVHD, while mice transplanted with SLP76pTYR transduced T cells survived for > 40 days post-HSCT and tumor challenge with minimal signs of GVHD and **(Fig. 9A-E)**.

**Figure 9.**
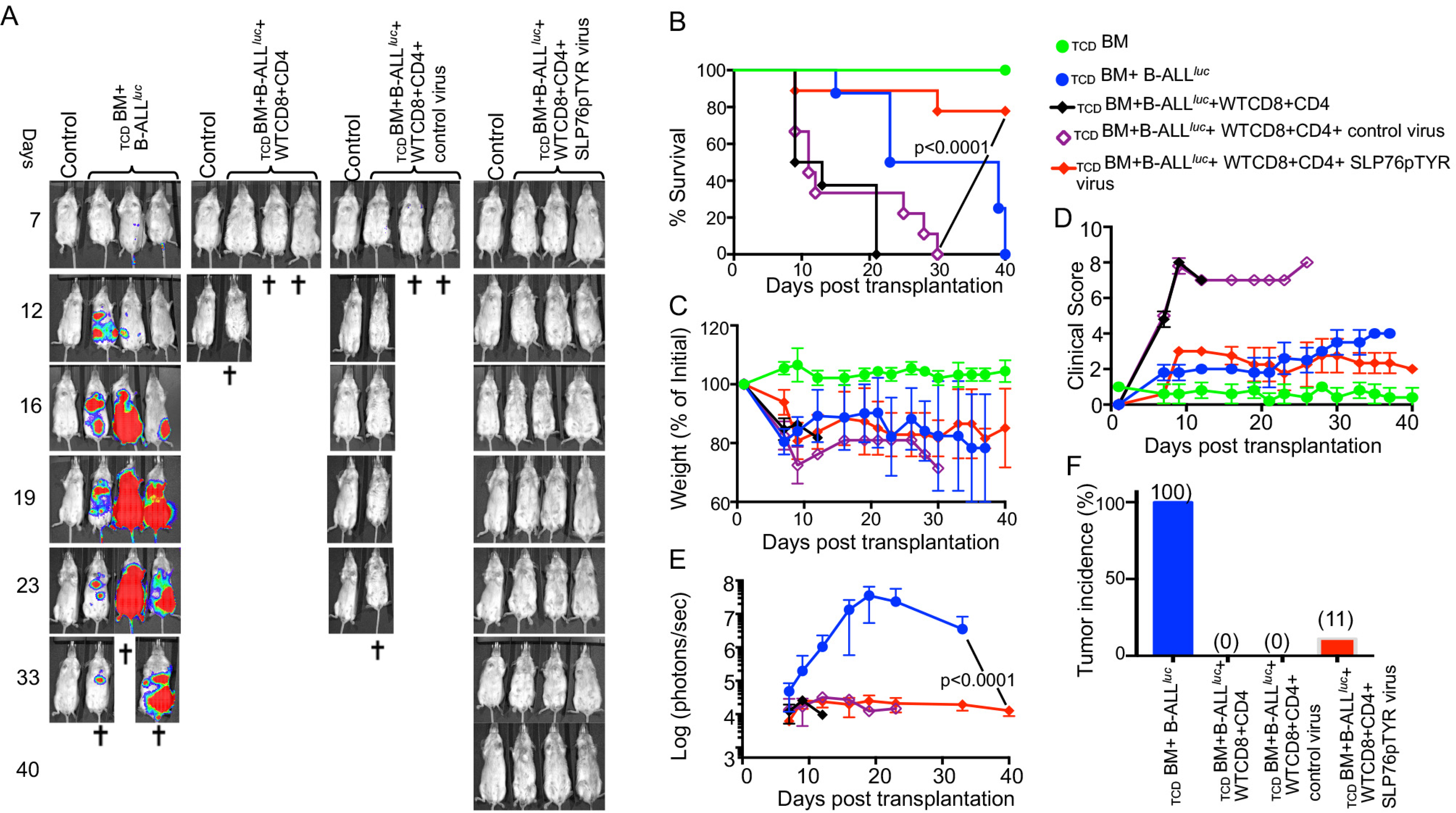
Inhibition of T cells by peptide SLP76pTYR allows tumor clearance without inducing GVHD. Purified WT CD8_+_ and CD4_+_ T cells were mixed (1X10_6_ each) at a 1:1 ratio, and transduced with viruses containing SLP76pTYR or empty vector, then transplanted along with 1X10_5_ B-ALL-*luc* cells into irradiated BALB/c mice. Host BALB/c mice were imaged using IVIS 3 times a week. **(A)** Group one received T cell depleted bone marrow alone (_TCD_BM). Group two received T cell depleted bone marrow along with 2X10_5_ B-ALL-*luc* cells, (_TCD_BM+B-ALL*_luc_*). The third group was transplanted with a 1:1 ratio of purified WT CD8_+_ and CD4_+_ T cells (1X10_6_ each) along with 2X10_5_ B-ALL-*luc* cells (_TCD_BM+B-ALL*_luc_*+WT CD8+CD4). Group four received a 1:1 ratio of purified CD8_+_ and CD4_+_ T cells (1X10_6_ each) transduced with control viruses along with 2X10_5_ B-ALL-*luc* cells (_TCD_BM+B-ALL*_luc_*+*Control* CD8+CD4). Group four received a 1:1 ratio of purified CD8_+_ and CD4_+_ T cells (1X10_6_ each) transduced with SLP76pTYR viruses along with 2X10_5_ B-ALL-*luc* cells (_TCD_BM+B-ALL*_luc_*+*Control* CD8+CD4) **(B)** The mice were monitored for survival, **(C)** body weight changes, and (**D)** clinical score for 40 days post BMT. For weight changes and clinical score, one representative of 2 independent experiments is shown (n = 3 mice/group for BM alone; n = 5 experimental mice/group for all three group). (**E**) Quantitated luciferase bioluminescence of tumor growth. Statistical analysis for survival and clinical score was performed using log-rank test and two-way ANOVA, respectively.

Altogether this data demonstrated that ITK signaling can separate GVHD from GVT. Inhibition of ITK by SLP76, specifically targeted ITK signaling, allows tumor clearance and minimizes development of GVHD. Finally, our novel peptide inhibitor of ITK is specific and has the potential to be used in a clinical setting for T cell-mediated disorders.

## Discussion

In this report, we demonstrate that the absence of the TCR-regulated kinase ITK significantly suppresses GVHD pathogenesis, while maintaining GVT in models of allo-HSCT. This is due to the effect of the ITK signaling pathway on regulation of Eomes, a critical transcription factor required for T cell cytotoxic function. Loss of ITK also altered expression of IRF4, and the JAK/STAT pathway components JAK1, JAK2 and STAT, which play critical roles in controlling cytokine expression^46, 47^. Transcriptome analysis by RNA sequencing also revealed that ITK signaling controls chemokine receptor expression during this process, which in turn affects the ability of donor T cells to migrate to GVHD target organs. Taken together, these data suggest that ITK could represent a potential target for the separation of GVHD and GVT responses after allo-HSCT.

The adaptor protein SLP76 plays an important role regulating T cell activation downstream of TCR by assembling a multimolecular signaling complex that includes ITK. The phosphorylation of SLP76 at Y145 leads to the activation and recruitment of ITK, which phosphorylates PLCγ1, leading to its activation, mobilization of calcium, and activation of the NFAT transcription factor^48^. Hence, T cells that carry a Y145F mutation of SLP76 fail to phosphorylate and activate PLCγ1 in response to TCR stimulation^10^. Although T cells expressing the SLP76 Y145F mutant exhibit signaling defects downstream of TCR stimulation, not all T cell functions are lost when ITK recruitment and activation is defective. For example, both *Itk_-/-_* and SLP76 Y145FKI T cells can clear acute LCMV infection^49, 50, 51^. The similarity in the ability of *Itk_-/-_* T cells, and the SLP76 Y145FKI T cells in being able to induce GVT without GVHD indicates that the SLP76/ITK pathway controls these functions. Both *Itk_-/-_* and SLP76 Y145FKI CD8_+_ T cells develop into IMP cells (CD122_+_ CD44_hi_ phenotype) in the thymus, and it is possible that such cells are responsible for the GVHD and GVT effects we observe. However, in experiments where WT T cells are able to develop into IMPs, we find that they retained the capacity to induce acute GVHD as well as clear tumor in GVT, suggesting a T cell-intrinsic function of ITK in promoting GVHD during allo-HSCT.

IMP cells express significantly higher Eomes expression compared to their WT non-IMP counterparts. However, we found that IMP CD8_+_ T cells are not responsible for distinguishing GVHD and GVT. Similarly, the cytotoxicity of *Itk_-/-_* CD8_+_ T cells is not dependent on the IMP. However, we found that expression of the transcription factor Eomes is required for the in vitro cytolytic function of CD8_+_ T cells that lack ITK/SLP76 signaling. Similarly, *in vivo,* Eomes is required for the GVT effects of T cells that lack ITK/SLP76 signaling in the allo-HSCT model. We did note that to our surprise, *Itk_-/-_* CD8_+_ T cells exhibit similar or higher *in vitro* cytotoxicity compared to WT CD8_+_ T cells. This may be due to the higher levels of perforin expressed by *Itk_-/-_* T cells compared to WT T cells. However, SLP76Y145FKI CD8_+_ T cells show slightly reduced cytotoxicity compared to WT cells. There may also be subtle differences in signaling between cell lacking *Itk* T cells, or carrying the SLP76Y145F mutant, that explains these differences.

Our data showed that *Itk_-/-_* donor CD8_+_ T cells exhibited reduced expression of chemokine receptors compared to WT counterparts. Moreover, the migration of *Itk_-/-_* donor T cells to target organs was also severely defective, reflecting the reduced expression of key chemokine receptors. The defective migration of *Itk_-/-_* CD8_+_ T cells likely contributes to the attenuation of GVHD, since these T cells could still display GVT effects against tumor cells that were injected intravenously and reside in secondary lymphoid organs. By contrast, WT but not *Itk_-/-_* CD8_+_ T cells were able to inhibit tumor growth when the tumor cells were injected subcutaneously. The compartmentalization of T cells to secondary lymphoid organs can be an effective strategy for preventing GVHD, while leaving GVT effects against hematologic malignancies intact. For example, the retention of T cells to secondary lymphoid organs by FTY720-mediated inhibition of S1P1 ameliorates GVHD while maintaining GVT effects^52, 53^. Similarly, inhibition of T cell migration to GVHD target organs by targeting the chemokine receptors CCR2 or CCR5 protects against GVHD-induced pathology^54, 55, 56^, which at least with CCR2 deficiency was shown to preserve the GVT effect. Importantly, in a clinical study, CCR5 blockade by a small molecule antagonist led to a reduction in GVHD with no significant difference in relapse rates, suggesting that blocking T cell migration to target tissues could reduce GVHD severity without compromising the beneficial GVT effect. In addition, the inhibition of CXCR3 ameliorates GVHD in allo-HSCT mice^57, 58, 59, 60^. Activated allo-reactive CD8_+_ T cells upregulate the expression of CX3CR1 and CXCR6 after allo-HSCT^61, 62^, and these receptors are important for the homing of CD8_+_ T cells to the liver and intestines. Thus, CXCR6 deficiency or blockade of the CXCR3 and CXCR6 ligands attenuates GVHD^60^, and importantly, the GVT effect is still maintained under these conditions^62^. Thus, blocking T cell migration by chemokine receptor blockade could be beneficial in the treatment of GVHD after allo-HSCT. Since activated *Itk_-/-_* CD8_+_ T cells displayed significantly reduced expression of chemokine receptors, the compartmentalization of CD8_+_ T cells to secondary lymphoid organs likely contributes to the preservation of GVT effects while severely attenuating GVHD.

Although suppression of TCR signaling can prevent GVHD, the complete suppression of T cell responses negates the beneficial GVT effect that is also provided by the same donor T cells after allo-HSCT^63, 64^. Thus, the fact that mice transplanted with *Itk_-/-_* T cells are able to mount GVT responses is an exciting feature. The preservation of the GVT response could have occurred for several reasons. First, the proliferation and cytotoxicity activity of *Itk_-/-_* CD8_+_ T cells is preserved compared to pro-inflammatory cytokine production. The manifestations and severity of GVHD are highly influenced by local cytokines, which, in turn, activate transcription factors and drive development toward cytokine storm. In addition, proinflammatory cytokines exert direct effects on GVHD target tissues^65, 66, 67^. Indeed, the presence of cytokine storm is considered one of the hallmarks of GVHD pathogenesis^26, 68^, and our data showed cytokine production was significantly reduced in mice that received *Itk_-/-_* or SLP76 Y145FKI T cells. We also confirmed that cytokine production is T cell-intrinsic while proliferation is T cell-extrinsic.

To explore the potential mechanism of this observed difference in cytokine and chemokine receptor expression between WT and *Itk_-/-_* donor T cells, we analyzed key transcription factors and pathways that may be involved in these processes. We found significant differences in expression of the transcription factor IRF4 and the JAK/STAT signaling pathways, which regulate the expression of key molecules required for the maintenance of T cell effector function, cytokine production, and chemokine receptor upregulation. Since IRF4 has been shown to play critical roles in modulating TCR signal strength and T cell function^11, 28, 69, 70^, it is likely that reduction in the activation of IRF4, and of the JAK/STAT pathway contribute to reduced cytokine expression, thus alleviating the cytokine storm in GVHD. In addition, the inability to migrate to target organs may also affect this process, and thus explain the inability of the *Itk_-/-_* donor T cells to induce GVHD.

We have also demonstrated proof of concept that specific targeting of the SLP76/ITK interactions can be achieved to potentially differentially modulate GVT and GVHD by pharmacological agents. Rather than directly inhibiting the activity of the kinase domain of ITK, which could result in complete blockage of all ITK kinase activity in T cells (and potentially non-specifically affect other tyrosine kinases), we developed a strategy to specifically disrupt the SLP76-pY145-mediated activation of ITK function in T cells. This strategy takes advantage of the SP76 pY145-mediated docking of ITK through its SH2 domain, given our findings that, like *Itk_-/-_* T cells, SLP76-Y145FKI mutant T cells can mediate tumor clearance through GVT, without inducing the unwanted GVHD effects. ITK interacts with SLP76 via its SH3 and SH2 domains onto the poly-proline motif and pY145 of SLP76 respectively. Thus, converting Y145 to F145 in SLP76 or preventing SH2 docking by our novel SLP76145pTYR peptide does not completely abolish the interaction between SLP76 and ITK, but significantly affects ITK kinase activity and results in severe defects in specific downstream signaling pathways^71, 72^. Therefore, we decided to target this specific interaction, which would retain signaling pathways that maintain GVT effects, but ameliorate GVHD. When we utilized SLP76145pTYR peptide to specifically target ITK signaling, we observed that this inhibitor is very specific in only inhibiting ITK signaling, without having significant effects on other tyrosine kinases nor apparent toxicity as determined by cell viability compared to nonspecific peptide or vehicle alone. Furthermore, SLP76145pTYR inhibition of ITK signaling also enhances Tregs i*n vitro*, confirming its ability to affect ITK signaling on T cell effector function. Our data demonstrated that TCR stimulation of primary human T cells isolated from PBMC in the presence of SLP76pTYR led to significantly reduced IFN-γ and TNF-α production, which was not observed when they were stimulated with PMA+Ionomycin in the presence of SLP76pTYR. Finally, donor T cells treated with SLP76pTYR, resulted in tumor clearance without inducing GVHD. Future combinatorial therapies involving our novel SLP76145pTYR peptide inhibitor and small molecule inhibitors, such as the BTK/ITK dual antagonist, Ibrutinib, can potentially be an effective strategy for enhancing GVT while avoiding GVHD. Ibrutinib is FDA-approved for the treatment of chronic lymphocytic leukemia^73^, and although its effects on GVT outcome has not been reported, Ibrutinib has recently been shown to protect against acute and chronic GVHD in mouse models of allogeneic BMT^74, 75^. Thus, combining more selective ITK inhibition using our SLP76Y145 peptide with Ibrutinib could be beneficial in the treatment of GVHD, while maintaining GVT effects after allo-HSCT.

## Materials and Methods

### Mice

SLP76 Y145FKI and SLP76 Y145FKI Eomes_flox/flox_ mice were generated as previously described and were a kind gift of Dr. Martha S. Jordan (University of Pennsylvania)_25_ *Itk_-/-_* mice as were described previously^76^. C57BL/6, C57BL/6.SJL (B6-SJL), ROSA26-pCAGGs-LSL-Luciferase, Thy1.1 (B6.PL-Thy1a/CyJ), CD45.1 (B6.SJL-Ptprc_a_ Pepc_b_/BoyJ) and BALB/c mice were purchased from the Charles River or Jackson Laboratory. Mice expressing Cre driven by the CMV promoter (CMV-Cre) were purchased from the Jackson Laboratory and crossed to ROSA26-pCAGGs-LSL-Luciferase mice (B6-luc). B6-luc mice were bred with SLP76Y145FKI or *Itk_-/-_* mice to create Y145F-luc mice and *Itk_-/-_luc*. *Itk_-/-_* and *Il4ra*_/-_ double knockout mice have been described^13^. Mice aged 8-12 weeks were used, and all experiments were performed with age and sex-matched mice. Animal maintenance and experimentation were performed in accordance with the rules and guidance set by the institutional animal care and use committees at SUNY Upstate Medical University and Cornell University.

### Reagents, cell lines, flow cytometry

Monoclonal antibodies were purchased from eBiosciences (San Diego, CA) or BD Biosciences (San Diego, CA). Antibodies used included anti-CD3, anti-CD28, anti-CD3-FITC, anti-CD8-FITC, anti-BrdU-APC, anti-IFN-γ-APC, anti-TNF-α-PE, anti-CD45.1-PerCPCy5.5, anti-CD122-APC, anti-CD44-Violet H, anti-Eomes-Blue A, anti-CD25-Violet H, anti-FoxP3-APC, anti-T-bet-Violet H, anti-CD4-Violet A, anti-CD45.1-Pacific Blue, anti-H-2K_d_-Pacific Blue. We used multiplex ELISA from Biolegend LEGEND plex and some kits were custom ordered to detect both mouse and human cytokines. Luciferin was purchased from Perkin Elmer (Waltham, MA) and Gold Bio (St Louis MO). Dead cells were excluded from analysis with LIVE/DEAD Fixable Aqua Dead Cell staining. Flow cytometry was performed by BD LSR-II or BD LSR Fortessa (BD Biosciences). Data were analyzed with FlowJo software (Tree Star, Ashland, OR).

For cell sorting, T cells were purified with either anti-CD8 or anti-CD4 magnetic beads using MACS columns (Miltenyi Biotec, Auburn, CA) prior to cell surface staining. FACS sorting was performed with a FACS Aria cell sorter (BD Biosciences). FACS-sorted populations were typically of > 95% purity. Antibodies against ITK, PLCγ1, ERK, IRF4, STAT3, JAK2, JAK1, GAPDH, β-Actin total and/or phospho proteins were purchased from Cell Signaling Technology (Danvers, MA). All cell culture reagents and chemicals were purchased from Invitrogen (Grand Island, NY) and Sigma-Aldrich (St. Louis, MO), unless otherwise specified. The A20 cell lines (American Type Culture Collection; Manassas, VA), and primary mice B-ALL blasts a primary cells ^16^ were transduced with luciferase, and cultured as described previously ^77^.

#### Statistics

All numerical data reported as means with standard deviation. Data are analyzed for significance with GraphPad Prism. Differences are determined using one-way or two-way ANOVA and Tukey’s multiple comparisons tests, or with a student’s t-test when necessary. P-values less than or equal to 0.05 are considered significant. All transplant experiments are done with N=5 mice per group, and repeated at least twice, according to power analyses. Mice are sex-matched, and age-matched as closely as possible.

### Allo-HSCT and GVT studies

Lethally irradiated BALB/c mice (800 cGy) were injected intravenously with 5×10_6_ T cell-depleted bone marrow (_TCD_BM) cells with or without either 1×10_6_ FACS-sorted CD8_+_, CD4_+_ T cells, or CD8/CD4 cells mixed at a 1:1 ratio. FACS-sorted total CD8_+_, total CD4_+_, or mixed CD8_+_ and CD8_+_ T cells from WT (C57Bl/6), *Itk_-/-_*, SLP76 Y145FKI or SLP76 Y145FKI Eomes_flox/flox_ mice either crossed with or without CD4cre were used. For GVT experiments, B-ALL primary blasts^16^ transduced with luciferase were cultured as described previously^77^ and 2×10_5_ luciferase-expressing primary B-ALL blasts were used. Mice were evaluated twice a week from the time of tumor injection for 70 days by bioluminescence imaging using the IVIS 200 Imaging System (Xenogen) as previously described^78^. Clinical presentation of the mice was assessed 2-3 times per week by a scoring system that sums changes in 5 clinical parameters: weight loss, posture, activity, fur texture, and skin integrity^17^. Mice were euthanized when they lost ≥ 30% of their initial body weight. Bone marrow cells from *Itk_-/-_* (CD45.1_+_) or C57Bl/6 (CD45.2_+_) mice were mixed at different ratios 1:1 (*Itk_-/-_*:WT), 1:2, 1:3, 1:4, and transplanted into lethally irradiated Thy1.1 mice. In some experiments, we used *Itk_-/-_* on a CD45.2 background and WT on a CD45.1 as indicated in the figure legends. Mice were bled through tail vein after 9 weeks to determine the presence of *Itk*_-/-_ and WT cells. In some experiments, *Itk*_-/-_ (CD45.1_+_) and WT (CD45.2) cells T cells were FACS-sorted from Thy1.1 hosts and then transplanted to irradiated BALB/c mice carrying tumor cells, along with T cell-depleted bone marrow as described above. This was followed by analysis of GVHD and GVT. In some experiments FACS-sorted CD8_+_ T cells from WT, SLP76 Y145FKI or *Itk_-/-_* B6 mice were mixed at a 1:1 ratio and injected into BALB/c mice (2×10_6_ CD8_+_ T cells total). For tissue imaging experiments, allo-HSCT was performed with 5×10_6_ WT T cell-depleted BM cells and 1×10_6_ FACS-sorted CD8_+_ T cells (from B6-luc, SLP76 Y145F-luc mice or *Itk*_-/-_*luc* mice) and bioluminescence imaging of tissues was performed as previously described ^79^. Briefly, 5 minutes after injection with luciferin (10μg/g body weight), selected tissues were prepared and imaged for 5 minutes. Imaging data were analyzed and quantified with Living Image Software (Xenogen) and Igor (Wave Metrics, Lake Oswego, OR).

### Cytokine production, cytotoxicity, and BrdU incorporation assays

On Day 7 post BM transplantation, serum and single cell suspensions of spleens were obtained. Serum IL33, IL-1α, IFN-γ, TNF-α and IL-17α content were determined by multiplex cytokine assays (Biolegend LEGEND plex). T cells were stimulated with anti-CD3/CD28 for 4-6 hours in the presence of brefeldin A (10μM) and stained intracellularly for cytokines (IFN-γ and TNF-α). Control cells were stimulated with PMA and ionomycin in the presence of brefeldin A. For detection of BrdU, mice were administered BrdU with an initial bolus of BrdU (2 mg per 200 μl intraperitoneally) and given drinking water containing BrdU (1 mg/ml) for 2 days. Spleen cells from these mice were resuspended in freezing media (90% FCS, 10% DMSO) and stored overnight at −80°C. Cells were subsequently thawed at room temperature and stained for BrdU.

For cytotoxicity assays, luciferase-expressing A20 cells were seeded in 96-well flat bottom plates at a concentration of 3×10_5_ cells/ml. D-firefly luciferin potassium salt (75 μg/ml; Caliper Hopkinton, MA) was added to each well and bioluminescence was measured with IVIS 200 Imaging System. Subsequently, ex vivo effector cells (MACS-sorted or FACS-sorted CD8_+_ T cells from bone marrow-transplanted mice) were added at 40:1, 30:1,20:1, 10:1, and 5:1 effector-to-target (E:T) ratios and incubated at 37°C for 4 hours. Bioluminescence in relative luciferase units (RLU) was then measured for 1 minute. Cells treated with 1% Nonidet P-40 was used as a measure of maximal killing. Target cells incubated without effector cells were used to measure spontaneous death. Triplicate wells were averaged and percent lysis was calculated from the data using the following equation: % specific lysis = 100 × (spontaneous death RLU– test RLU)/(spontaneous death RLU– maximal killing RLU).

### Migration assays

Lethally irradiated BALB/c mice were injected intravenously with 5×10_6_ WT T cell-depleted bone marrow (_TCD_BM) from B6.PL-*Thy1_a_*/CyJ and FACS-sorted CD8_+_ T cells from B6.SJL and *Itk_-/-_* or SLP76 Y145FKI mice mixed at a 1:1 ratio. Seven days post transplantation, the mice were sacrificed and lymphocytes from the liver, small intestine, spleen, and skin-draining lymph nodes were isolated. Livers were perfused with PBS, dissociated, and filtered with a 70μm filter. The small intestines were washed in media, shaken in strip buffer at 37°C for 30 minutes to remove the epithelial cells, and then washed, before digesting with collagenase D (100 mg/ml) and DNase (1mg/ml) for 30 minutes in 37°C, and followed by filtering with a 70 μm filter. Lymphocytes from the liver and intestines were further enriched using a 40% Percoll gradient. The cells were analyzed for H2K_b_, CD45.1_+_ and CD45.2_+_ CD8_+_ T cells by flow cytometry, but we excluded any bone marrow-derived T cells (Thy1.1_+_).

### RNA sequencing

CD8_+_ T cells from WT C5Bl/6 or *Itk_-/-_* mice were MACS purified and FACS sorted, and 2×10_6_ FACS sorted CD8_+_ T cells were transplanted into BALB/c mice, along with _TCD_BM as described above. Seven days post transplantation, donor cells were purified from spleen (Spl) and small intestine (Gut). Samples were submitted to SUNY Upstate Medical University Sequencing core facility for RNA sequencing. We were unable to sort enough donor T cells from small intestine of the recipient mice that received *Itk_-/-_* T cells. Therefore, we generated RNA sequencing data from five groups: WT-Pre and *Itk_-/-_*-Pre cells prior to transplantation; and WT-Gut, WT-Spleen, and *Itk_-/-_*Spleen using cells isolated from 7 days post transplantation. Copy numbers were further analyzed in Gene Spring for normalization, quality control, correlation, principal component analysis and gene differential expression. The sequencing data is deposited in (https://www.ncbi.nlm.nih.gov/geo/)

### Western blotting

Cells were lysed in freshly prepared lysis buffer (10 mM Tris, pH 8, 150 mM NaCl, 1 mM EDTA, 1% Nonidet-P40, 0.5% Deoxycholate, 0.1% SDS, complete Protease Inhibitor Cocktail [Roche, Palo Alto, CA], and 500 μM PMSF) and centrifuged for 10 minutes at 4°C. Aliquots containing 70 μg protein were separated on an 12-18% denaturing polyacrylamide gel and transferred to nitrocellulose membranes for immunoblot analysis using Abs specific to IRF4, JAK1, JAK2, ERK, PLCγ1, ITK, mTOR, AKT, P13K and STAT3 total and/or phospho proteins.

### qPCR assay

To confirm the differences observed in RNA sequencing, pre- and post-transplanted donor T cells were FACS sorted from recipient mice on H2K_b_ markers, and total RNA was isolated from T cells using the RNeasy kit Qiagen (Germantown, MD). cDNA was made from total RNA using cDNA synthesis kit (Invitrogen). qRT-PCR assay was performed with a premade customized plate (Fisher Scientific, Hampton, NH).

### SLP76145pTYR Peptide

To generate a molecule that specifically inhibist the interaction between pY145 of SLP76 and the SH2 domain of ITK, we designed a peptide based on the amino acid sequence of SLP76 from N132 to A155, which contains a phosphorylated tyrosine residue at Y145 (**Fig. 8**). To ensure that our peptide easily enters cells and that its cellular localization can be monitored, we incorporated a C-terminal TAT-peptide and an N-terminal fluorescent FITC dye respectively and named it SLP76145pTYR peptide (**Fig. 8**). Both SLP76145pTYR peptide (FITC Dye) - 132NEEEEAPVEDDAD**pY**EPPPSNDEEA155-(GRKKRRQRRRPQ) and non-specific (FITC Dye)-IIMTTTTNNKKSSRRVVVVAAAADD-(GRKKRRQRRRPQ) control peptide, were synthesized by Genscript Inc (Piscataway, NJ), and initially dissolved in 3% ammonia water to a final concentration of 10μg/μL and then further diluted into PBS media to a final concentration of 1μg/μL. Fresh splenocytes were isolated from WT mice, and T cells were generated from splenocytes as previously described^80^. Briefly, T cells were isolated from splenocytes using Miltenyi beads, then cultured in complete RPMI media (3×10_6_ cells/mL) plated on anti-CD3 (145-2C11; BD Pharmingen, San Diego, CA) antibody-coated tissue culture plates. T cells were incubated with SLP76145pTYR peptide or vehicle alone at different concentrations ranging from 10ng/ml to 1μg/ml in the presence of 4μg/ml of polybrene (Sigma-Aldridge, St. Louis, MO). Cells were spun for 60 minutes at 200 RPM and cultured for 24-48 hours 24 to 48 hours. Cells were examined for the presence of FITC by microscopy using a Leica DMi8 microscope equipped with an infinity total internal reflection fluorescence (TIRF) and DIC modules, a Lumencor SOLA SE II light box, a 150 mW 488 (GFP) laser and filter cube, a 100x/1.47 NA objective, and an Andor iXon Life 897 EMCCD camera. FITC expression was confirmed by flow cytometry as well Cells were lysed and used in Western blots.

### Toxicity assay

Yac1 and B-ALL cells line were transduced with luciferase as described ^81^. Luciferase-expressing tumor cells were placed in 96–well round bottom plates at a concentration of 3×10_5_ cells/ml in triplicates, given D-firefly luciferin potassium salt (75 µg/ml; Caliper Hopkinton, MA), and luciferase activity measured with a luminometer. This was done to establish the Bioluminescence Imaging (BLI) baseline readings before the occurrence of any cell death and to ensure equal distribution of target cells among wells. Subsequently, SLP76pTYR or vehicle were added at 1μg/well in 100μl of media and serial dilutions performed, followed by incubation at 37°C. BLI was then measured for 60 seconds with a luminometer (Packard Fusion Universal Microplate Analyzer, Model A153600) as relative light units (RLU). Triplicate wells were averaged and data presented as photons per second per well.

### Human Patient Samples

T cells were isolated from Peripheral Blood Mononuclear Cells (PBMC) of GVHD patients or normal healthy donors as previously described ^82^. Briefly, mononuclear cells were isolated from GVHD patient samples by Ficoll-Hypaque density centrifugation and separated by positive selection with CD8 and CD4 MACS Microbeads. The final product was resuspended at 2×10_6_ cells/ml in media and stimulated with OKT-3 (1 μg/ml; Ortho Bio Tech), anti-human CD28, in the presence SLP76pTYR or vehicle for 24 hours. T cell lysates were used in western blot analysis. We also isolated plasma from GVHD patients and healthy donors and performed cytokine ELISAs on these plasma samples using multiplex ELISA kits (Biolegend, San Diego, CA).

#### Transducing primary T cells with SLP76pTYR

To generate viruses that specifically express SLP76pTYR. SLP76pTYR was cloned as fusion protein with pCherry ordered through 14

Integrated DNA Technology (IDT). The insert was cloned into pQCX-I-X retroviral vector between MLU1 and Xho1 restriction sites, and the expression of insert confirmed by digestion and sequencing. To produce retroviral supernatants, Phoenix packaging cells were plated in 175 cm_2_ flasks, and transfected with 20 μg of vector using Lipofectamine 2000 reagent (Invitrogen, Carlsbad, CA), according to the manufacturer’s protocol. The medium was changed after 8–12 hours, and viral supernatants were harvested after 24–36 hours. Concentrated viral supernatants were re-suspended in IMDM media (Invitrogen) and used to transduce primary T cells in the presence of polybrene (10 μg/ml) and protamine sulfate (10 μg/ml) to enhance transduction efficiency. T cells transduced with viruses containing either SLP76pTYR or empty plasmid were injected into mice.

## Acknowledgements

We thank all members of the Karimi and August laboratories for helpful discussions. This research was funded in part by a grant from the National Blood Foundation Scholar Award to (MK) and the National Institutes of Health (NIH LRP #L6 MD0010106 and AI130182 to MK, AI120701 and AI126814 to AA, R35ES028244 to AA and Gary Perdew, AI129422 to AA and WH, and AI146226 and GM130555-sub6610 to WH).

## Author contributions

MM, WH, AS, QY, SD, JR, AB and MK performed experiments; WT, YC, JP and TG provided valuable reagents; RH assisted with data analysis, experimental design, and scientific discussion; and WH, QY, AA, AB and MK designed experiments, analyzed the data, and wrote the manuscript.

## Supplementary Figure Legends

**Supplementary Figure 1. CD4_+_ T cells from *Itk_-/-_* and SLP76 Y145F mice exhibit attenuated induction of GVHD compared to WT T cells. (A, D)** 1X10_6_ purified WT or *Itk_-/-_* CD4_+_ T cells (A) or WT or SLP76 Y154FKI CD4_+_ T cells (D) were transplanted into irradiated BALB/c mice. The mice were monitored for survival, **(B, E)** changes in body weight, **(C, F)** clinical score for 70 days post BMT. For weight changes and clinical score, one representative of 2 independent experiments is shown (n = 3 mice/group for BM alone; n = 5 experimental mice/group for all three groups). The and p vales are presented. Two-way ANOVA and Student’s *t* test were used for statistical analysis.

**Supplementary Figure 2. Disruption of ITK SLP76 Y145 signaling allows tumor clearance without inducing GVHD.** 1X10_6_ purified WT or SLP76 Y145FKI CD8_+_ T cells, and 1X10_6_ purified CD4_+_ T cells were mixed at a 1:1 ratio, and transplanted along with 2X10_5_ A20-*luc* cells transplanted into irradiated BALB/c mice. Host BALB/c mice were imaged using IVIS 3 times a week. **(A)** Group one received T cell depleted bone marrow alone (_TCD_BM). Group two received T cell depleted bone marrow along with 2X10_5_ A20-*luc* cells, (_TCD_BM+B-ALL*_luc_*). The third group was transplanted with 1X10_6_ purified WT CD8_+_ and 1X10_6_ CD4_+_ T cells (1:1 ratio) along with 2X10_5_ A20-*luc* cells (_TCD_BM+B-ALL*_luc_*+WT CD8+CD4). Group four received 1X10_6_ purified SLP76 Y145FKI CD8_+_ and 1X10_6_ CD4_+_ T cells (1:1 ratio) along with 2X10_5_ B-ALL-*luc* cells (_TCD_BM+A20*_luc_*+*Itk_-/-_* CD8+CD4). **(B)** The mice were monitored for survival, **(C)** body weight changes, and (**D)** clinical score for 65 days post BMT. For weight changes and clinical score, one representative of 2 independent experiments is shown (n = 3 mice/group for BM alone; n = 5 experimental mice/group for all three group). (**E**) Quantitated luciferase bioluminescence of tumor growth. Statistical analysis for survival and clinical score was performed using log-rank test and two-way ANOVA, respectively. *Note: Control mouse is naïve mice used negative control for BLI*.

**Supplementary Figure 4. Eomes expression in T cells from *Itk/Il4ra* DKO and SLP76 Y145FKI Eomes_flox/flox_ mice. (A)** Purified T cells from WT and *Itk/Il4ra* DKO mice were isolated and transplanted into irradiated BALB/c mice. At day 7 donor T cells were gated for expression of H-2K_b_ and CD45.2, and intracellular expression of Eomes pre and post-transplantation was analyzed. **(B, C)** Purified T cells from SLP76 Y145FKI Eomes_flox/flox_ mice with or without CD4cre were transplanted into irradiated BALB/c mice. Pre- **(B**) and 7 days post-transplantation **(C)** of T cells from WT or SLP76 Y145FKI Eomes_flox/flox_ mice with or without CD4cre, T cells were examined for Eomes expression.

**Supplementary Figure 4. T cells from *Itk_-/-_* mice are capable of cytokine production.** Purified WT and *Itk_-/-_* T cells were transplanted into irradiated BALB/c mice. At day 7 donor T cells were gated for expression of H-2K_b_, and CD45.2 and analyzed for intracellular expression of IFNγ and TNF-a following *ex vivo* stimulation with PMA/ionomycin. Data from several experiments were combined and statistical analysis performed using two-way ANOVA and Student’s *t* test, with p values presented

**Supplementary Figure 5. 10n and CTA056 ITK inhibitors are not specific. (A)** WT T cells were cultured with either 10n or vehicle, then lysed post incubation and lysates were western blotted for pITK, pPLCγ1 and pERK, pAKT, and pMTOR. (**B**) Western blots from three experiments were quantitated and normalized to βActin. **(C)** WT T cells were cultured with commercially available ITK inhibitor CTA056 or vehicle, and western blotted for pITK, pPLCγ1 and pERK, pAKT, and pMTOR. **(D)** Western blots from three experiments were quantitated and normalized to βActin. **(E)** T cells from primary human PBMCs were isolated and cultured with either 10n or vehicle, the cells were lysed post incubation and lysates were western blotted for pPLCγ1 pAKT, pMTOR and pERK. **(F)** Western blots from three experiments were quantitated and normalized to βActin. Two-way ANOVA and Student’s *t* test were using for statistical analysis.

**Supplementary Figure 6. SLP76pTYR peptide is not toxic to cells.** Yac1 and B-ALL-*luc* cells were cultured in 96 well plates in 3 replicates in the presence of vehicle only or increasing concentrations of SLP76pTYR as indicated. Cell were examined at 0 hrs., 2 hrs., 4 hrs. and 6 hrs. Two-way ANOVA and Student’s *t* test were used for statistical analysis.

